# Evolutionary trajectory of TRPM2 channel activation by adenosine diphosphate ribose and calcium

**DOI:** 10.1101/2024.03.18.584035

**Authors:** Cheng Ma, Yanping Luo, Congyi Zhang, Cheng Cheng, Ning Hua, Xiaocao Liu, Jianan Wu, Luying Qin, Peilin Yu, Jianhong Luo, Fan Yang, Lin-Hua Jiang, Guojie Zhang, Wei Yang

## Abstract

Ion channel activation upon ligand gating triggers a myriad of biological events and, therefore, evolution of ligand gating mechanism is of fundamental importance. TRPM2, a typical ancient ion channel, is activated by adenosine diphosphate ribose (ADPR) and calcium and its activation has evolved from a simple mode in invertebrates to a more complex one in vertebrates, but the evolutionary process is still unknown. Molecular evolutionary analysis of TRPM2s from more than 280 different animal species has revealed that, the C-terminal NUDT9-H domain has evolved from an enzyme to a ligand binding site for activation, while the N-terminal MHR domain maintains a conserved ligand binding site. Calcium gating pattern has also evolved, from one Ca^2+^-binding site as in sea anemones to three sites as in human. Importantly, we identified a new group represented by olTRPM2, which has a novel gating mode and fills the missing link of the channel gating evolution. We conclude that the TRPM2 ligand binding or activation mode evolved through at least three identifiable stages in the past billion years from simple to complicated and coordinated. Such findings benefit the evolutionary investigations of other channels and proteins.

## 1. Introduction

Modification of protein functions by amino acid substitution plays a key role during species evolution and adaptation. The type, function and regulation of large biological molecules such as proteins have become increasingly more sophisticated from invertebrates to vertebrates. There are few detailed studies of the evolutionary trajectories of proteins, especially for those having experimental evidence besides analytical supports and for those investigated the molecular evolution in a long time scale. Recent researches have highlighted the molecular evolutionary trajectories and the underlying mechanisms of proteins that play a key role in brain development, oxygen delivery, ion transportation, and other vital biological processes. For example, a study examining haemoglobin evolution found that very few genetic changes might have driven the evolution of haemoglobin complex structure and function from its dimeric ancestral form to the current heterotetrameric form [1]. The transition of a protein from an ancestral form to the current structure is often accompanied with gain or loss of function or alterations in oligomeric states [1] or substrate specificities [2, 3]. The evolution of protein function could be driven by genetic changes [1, 4, 5], accelerated by cryptic genetic variations [6] and entrenched by epitasis events [2, 3, 7–10]. Understanding how the amino acid composition and structure of a protein evolved to govern its function can provide new insights into its function.

Ion channels as ancient membrane proteins mediate ion influx or efflux across the cellular membranes, and play an essential role in regulating a multitude of physiological processes such as cell volume, neuronal excitability, muscle contraction and nutrient transportation [11]. A great number of diseases arise due to disruption of the normal ion channel function, and are also thought to play a fundamental role in the evolution of animals [12]. Transient receptor potential melastatin 2 (TRPM2) forms a Ca^2+^-permeable non-selective cation channel that plays an important role in mediating oxidative stress related diseases [13], neutrophil chemotaxis [14], insulin secretion [15] and regulation of body temperature [16]. TRPM2 represents one of the most ancient ion channels, as it is found in early metazoans such as unicellular protists (*Monosiga brevicollis*) and sea anemones (*Nematostella vectensis*) [17], and in almost all metazoan species including invertebrates and vertebrates. The long evolutionary history of TRPM2 makes it a good candidate for inferring how the evolutionary changes on this protein have modulated the channel activation mode during a long period of time and contributed to species adaptation.

The TRPM2 channel is a tetrameric complex, and each subunit is composed of an intracellular N-terminus containing four TRPM homology regions (MHR1-4), a transmembrane domain (TMD) consisting of six transmembrane segments (S1-S6), and an intracellular C-terminus containing a coiled-coil domain and a NUDT9 homology (NUDT9-H) domain [18]. It can be activated by intracellular ADP-ribose (ADPR), a product of oxidative stress and DNA damage repair [19–23], that plays an essential role in cellular signaling under many physiological and pathological conditions. Recent studies have disclosed that the ADPR binding and gating mode differs dramatically in TRPM2 channels from different species. The TRPM2 channel from *Nematostella vectensis* (nvTRPM2) only requires ADPR binding to the N-terminal MRH1/2 domains for channel activation [24]. The C-terminal NUDT9-H domain functions as a pyrophosphatase that can hydrolyze ADPR to AMP and ribose-5-phosphate and is dispensable for channel activation [24, 25]. As for the TRPM2 channel from human (hsTRPM2), both the N-terminal MRH1/2 domains and the C-terminal NUDT9-H domain are critical for ADPR binding and channel gating [26, 27]. Furthermore, activation of the hsTRPM2 channel requires intracellular Ca^2+^ [28, 29], a ubiquitous second messenger that mediates all kinds of body and cell functions. Studies have identified three major domains that are involved in the hsTRPM2 channel activation by Ca^2+^ [25, 30–33], including an IQ-like motif [31] and an EF-loop motif [33] in the N-terminus, and the Ca^2+^-binding pocket formed by the S2-S3 helix [25, 30, 32]. There is no information regarding the role of intracellular Ca^2+^ in the activation of TRPM2 channels in other species. Furthermore, there are no systematical studies of the evolutionary trajectories of the gating modes of TRPM2 channels across different species, and the molecular mechanisms by which such differences in the gating modes evolved.

In this study, we provide analytical and experimental evidences to show that the mode of TRPM2 channel activation by ADPR and Ca^2+^ evolved from a relatively simple mode to a complicated and highly coordinated one through at least three identifiable stages. We systematically analyzed TRPM2 protein sequences from more than 280 species from invertebrates to vertebrates, and illustrated the sequence evolution in the domains and motifs engaged in the TRPM2 channel activation by ADPR and Ca^2+^. We have identified a group of TRPM2 proteins, represented by *Oryzias latipe* TRPM2 existing in the majority of fishes, have a new gating mode that represents the missing nexus in the evolution of ligand binding and gating. Therefore, the mode of TRPM2 channel activation by ADPR and Ca^2+^ evolved from a relatively simple mode in TRPM2s from Cnidaria, such as the nvTRPM2, to an intermediate one in TRPM2 channels from fish, represented by the olTRPM2, and to a more complicated and coordinated one in the hsTRPM2. Important permissive changes at the position corresponding to Y1349 in the hsTRPM2 indicates the requirement of the NUDT9-H domain for channel activation by ADPR, and this activation mode may have resulted from the environmental oxygen level as the selective pressure.

## 2. Materials and Methods

### 2.1 Cell culture

Human embryonic kidney 293T cells (HEK293T, ATCC (CRL-3216)) were maintained in Dulbecco’s modified Eagle’s medium (DMEM, Gibco) supplemented with 10% (v/v) fetal bovine serum (Gibco), 100 units/mL penicillin and 100 mg/mL streptomycin at 37°C with 5% CO_2_.

### 2.2 Data collection

The collection of TRPM2 and TRPM2-like protein sequences was assembled by performing Blast mainly in the Uniprot database (https://www.uniprot.org/), using hsTRPM2 protein sequence (O94759) as an input. This collection was expanded to include several invertebrate TRPM2 or TRPM2-like protein sequences [34] sourced from the NCBI and genomic database JGI (http://www.jgi.doe.gov/).

To improve the quality of the initial sequence collection, we performed sequence alignments using Clustal Omega [35]. Based on initial alignments, we removed duplicates and sequences lacking the C-terminal NUDT9-H domain or shorter than one third of the hsTRRM2 sequence, namely, without the majority of the key elements and domains. We retained the longest isoform for each species in order to maximize the available phylogenetic information. After several rounds, a dataset containing 89 TRPM2/TRPM2-like protein sequences was assembled to cover representative species from invertebrates, *Branchiostoma*, fish, amphibians, reptiles, birds and mammals.

To strengthen the reliability of data analysis and the conclusions drawn from phylogenetic interpretations, a second dataset, which is composed of 289 TRPM2 sequences with the majority of them collected from the NCBI database following the same logic described above, was analyzed independently of the first dataset.

### 2.3 Phylogenetic trees and divergence time analyses

Phylogenetic trees were reconstructed for datasets containing TRPM2 and TRPM2-like protein sequences using MEGA version 10.0.4. We firstly determined that the Jones-Taylor-Thornton (JTT) model with gamma distributed with variant sites (G) [36] was the best model. The phylogenetic trees were constructed using the Maximum Likelihood (ML) method based on the JTT matrix-based model [36]. The phylogenetic trees were inferred using the bootstrap method with 1000 replicates taken to represent the evolutionary history of our datasets [37]. The divergence times of the phylogenetic tree was analyzed through the “clock” module of MEGA version 10.0.4. The most ancient and divergent sequence of *Salpingoeca rosetta* was used as the outgroup.

We surveyed the literatures of the past molecular clock analyses (Supplementary references for section 2.3) from the TimeTree database [38] to assemble our calibration sets, which are (1) Primitive: *Branchiostoma floridae* and *Strongylocentrotus purpuratus*, the median time deducted based on the cited publications is 627 MYA [Sup.1-10] and this median divergence time is comparable to the most recently published data (600 MYA) [Sup.10]; (2) Cartilaginous fishes and Amphibian: *Callorhinchus milii* and *Xenopus laevis*, the deducted median divergence time 465 MYA [Sup.8, 9, 11-22] is close to the newest data (440.8 MYA) [Sup.22]; (3) Amphibian and Reptile: *Xenopus laevis* and *Chelonia mydas*, the median divergence time 351.7 MYA [Sup.9, 11, 14-18, 23-36] is comparable to the newest publication (349.9 MYA) [Sup.36]; (4) Among Fishes: *Oryzias latipes* and *Danio rerio*, the median divergence time 224 MYA [Sup.10, 15-17, 19-22, 29, 30, 37-57] agrees with the most recently published data (224 MYA) [Sup.22]; (5) Birds and Reptile: *Columba livia* vs *Chelonia mydas*, the median divergence time 262 MYA [Sup.10, 18, 20, 21, 29-34, 36, 58-72] is comparable to the recently published one (244.1 MYA) [Sup.36]; and (6) Among Primates: *Cebus capucinus imitator* and *Homo sapiens*, the median divergence time 42.9 MYA [Sup.25, 35, 73-100] agrees with the newest one (42.4 MYA) [Sup.100].

### 2.4 Constructs

The plasmids containing the full-length hsTRPM2 cDNA sequence (https://www.ncbi.nlm.nih.gov/gene, protein accession number: NP_003298.1) and mouse TRPM2 cDNA sequence (protein accession number is Q91YD4) in pcDNA3 vector were from Jiang’s lab (Leeds University, United Kingdom). The full-length *Nematostella vectensis, Oryzias latipes* and *Danio rerio* TRPM2 cDNA sequences (accession number of the corresponding protein: jgi.Nemve1.248535, H2MNZ1 and S5TZ89, respectively) were custom-synthesized by Union Gene Company and subcloned into pcDNA3.1 vector.

### 2.5 Cell preparations for electrophysiology

HEK293T cells were grown at 37°C with 5% CO_2_ in Dulbecco’s modified Eagle’s medium (DMEM, Gibco) supplemented with 10% (v/v) fetal bovine serum, 100 units/mL penicillin and 100 mg/mL streptomycin. One day before transfection, cells were re-suspended in DMEM and seeded onto glass coverslips in 35-mm dishes at 10-40% confluency. On the next day, cells were co-transfected using Lipofectamine 3000 Transfection Reagent (Invitrogen) with plasmids expressing wildtype (WT) or mutant TRPM2 channel and pEGFP-N1 (Invitrogen) that expresses enhanced green fluorescent protein (GFP) to permit visual identification of transfected cells at a ratio of 2:1. Cells were used for electrophysiological experiments 24 h after transfection.

### 2.6 Electrophysiology

Whole-cell patch clamp recordings were made at room temperature, using an Axopatch 700B amplifier (Molecular Devices). The extracellular solution contained 147 mM NaCl, 2 mM CaCl_2_, 10 mM HEPES and 13 mM glucose, pH 7.4. The intracellular solution contained 147 mM KCl, 0.05 mM EGTA, 10 mM HEPES, and ADPR or CaCl_2_ at indicated concentrations, pH7.4. The currents were recorded by applications of a 500 ms voltage ramp from -100 to +100 mV every 5 s, with a holding potential of -40 mV. Cells transfected with plasmids were placed in a recording chamber and continuously perfused with extracellular solution. After the patch pipette formed a seal of ≥1 GΩ with the cells, three sweeps were recorded as the baseline. Subsequently, negative pressure was applied to establish a whole-cell recording configuration. Once the current reached a steady state, 20 µM N-(pamylcinnamoyl) anthranilic acid (ACA) (MCE) was applied via extracellular perfusion to block any substantial residual currents at the end of each recording. Only the recordings showing complete inhibition of currents by ACA were used. The perfusion and solution switching were achieved using a gravity-driven system at a rate of 400 ms per tube. The tubes were positioned directly towards the cells, ensuring rapid solution exchange within a short period of time. Glass pipettes with a resistance of 3–5 MΩ were used. Data were acquired at 10 kHz and filtered offline during data analysis. Change of the extracellular solutions was performed using a RSC-160 system (Bio-Logic Science Instruments) with a solution change time of ∼400 ms.

### 2.7 Surface expression of WT and mutant TRPM2 channels

New plasmids expressing WT and mutant TRPM2 from different species carrying GFP at the C-terminus and a FLAG epitope inserted into the extracellular S3-S4 loop were constructed for assessing the mutational effects on surface expression detection by flow cytometry. HEK293T cells were transfected with the WT or mutant plasmid for 24 h. Cells were incubated in a solution (147 mM NaCl, 2 mM CaCl2, 10 mM HEPES, and 13 mM glucose, pH 7.4) containing an anti-FLAG antibody produced in mouse (F1804, Sigma) and subsequently with a red fluorescent fluorophore-conjugated secondary anti-mouse IgG antibody (130522, EARTH) to label TRPM2 proteins on the plasma membrane. Cells were washed three times with a buffer containing 147 mM NaCl, 2 mM CaCl_2_, 10 mM HEPES, and 13 mM glucose, pH 7.4, before being collected for flow cytometry. Red fluorescence (PE-Taxas Red channel, ex488nm/em615nm) is contributed by the TRPM2 proteins expressed on the cell surface, while green fluorescence (FITC channel, ex488nm/em530nm) reflects the total protein expression. The proportion of cells with red fluorescence to cells with green fluorescence indicates the level/ratio of TRPM2 surface expression.

### 2.8 Data analysis and statistics

Prism 8 software (GraphPad Software) was used for statistical analyses. The data are presented in the text and figures as mean ± SEM value. The currents mediated by the mutant channels were normalized with the mean maximal currents by the WT channels recorded on the same day and therefore referred to as the relative currents. P < 0.05 was considered to be statistically significant.

## 3. Results

### 3.1 Evolution of TRPM2 on sequence level: phylogenetic and molecular evolutionary analysis of TRPM2 protein sequences

We performed phylogenetic analysis of TRPM2 protein sequences from 89 species encompassing a diverse range of organisms (Supplementary Table S1). They include *Salpingoeca rosetta*, representing the “oldest” unicellular organisms, *Trichoplax adhaerens, Nematostella vectensis*, *Capitella teleta* and *Strongylocentrotus purpuratus*, each representing a different invertebrate phylum, *Branchiostoma floridae*, a “living-fossil” representing Cephalochordata, and species from key vertebrate groups. The unrooted phylogenetic tree of TRPM2 proteins (supplementary Figure S1) well reflects the species phylogeny. We also collected another dataset containing 289 TRPM2 protein sequences and performed the same analysis independently. The results from such analysis agree well with our observations (Figure 1A and Table S4).

**Figure 1.**
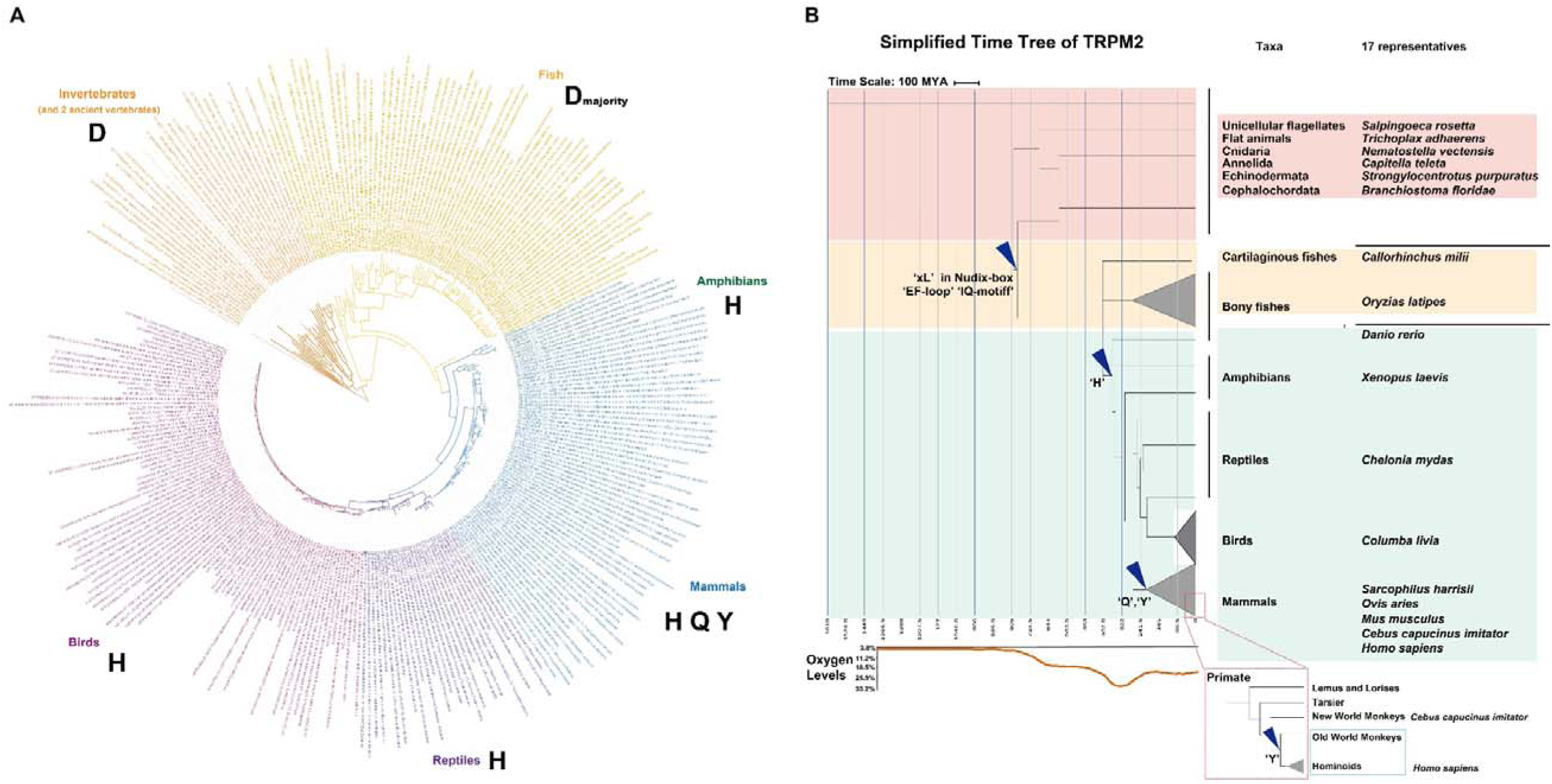
Maximum likelihood tree and simplified tree indicating the divergence times of TRPM2 channels. (1) TRPM2 and TRPM2-like protein sequences of 289 species, from Unicellular flagellates, Flat animals, Jellyfish/anemones, Worms, Sea stars/urchins, Lancelets, Cartilaginous fishes, Bony fishes, Amphibians, Reptiles, Birds and Mammals. The amino acids corresponding to Y1349 in hsTRPM2 were labeled for each taxa (datasets information could refer to Table S4). (A) (B) A simplified tree indicating the divergence times of TRPM2s (original tree see Figure S2) shown with taxa and 17 corresponding representative species listed on its right. The curves of the oxygen level related to time were shown below the tree.

In order to deduce the divergence time, we constructed the phylogenetic trees (Figure 1B and supplementary Figure S2) with the most ancient and divergent sequence of *Salpingoeca rosetta* as the outgroup using the ML methods and the JTT+G model. The divergence times of phylogenetic trees were further calibrated as detailed in the Methods and Materials section. To note, all the observations are based on the analysis of TRPM2 protein sequences from the full dataset and, for clarity, part of the alignments of 17 representatives are shown in figures.

### 3.2 Evolution of TRPM2 on sequence level: evolutionary changes in the domains engaged for ADPR binding and activation

To reveal the sequential evolutionary process of TRPM2 proteins, we first detected the present/absent pattern of domains that are required for ADPR binding in the N-terminal MHR1/2 domains and the C-terminal NUDT9-H domain [25]. We identified amino acid residues that are potentially involved in forming the N-terminal ADPR-binding pocket or near the pocket by calculating their distances to the bound ADPR molecule based on the atomic structure of hsTRPM2 (6PUS) (Figure 2A and B, and Figure 3A). The conservation scores of identified residues were calculated based on the 89 TRPM2 sequences from Consurf [39] and are generally lower than -0.728, with the conservation color scores over 8. The residues forming the N-terminal ADPR-binding site are present even in the ancient TRPM2 or TRPM2-like proteins in *Salpingoeca rosetta*. The residues forming the ADPR-binding pocket in the C-terminal NUDT9-H domain are also highly conserved. Interestingly, the residue corresponding to Y1349 in hsTRPM2 shows a clear taxa-specific pattern (Figure 1, Figure 2A, 2C and supplementary Figure S1, S7). Of all 29 invertebrate species, the TRPM2 sequences all possess aspartic acid (D) at this position (Figure 1A and supplementary Table S4). This amino acid type was preserved in the majority of fish species, with 49 of the 69 fish TRPM2 sequences containing D and 12 having glutamic acid (E) (Figure 1A and supplementary Table S4). Other fish species independently evolved with histamine (H) in this position, which also appeared in common ancestors of tetrapods (Figure 1B). The H amino acid type was then present during the diversification of amphibian, reptiles and birds. The mammalian group possesses the most diverse pattern and evolved with glutamine (Q) at this position in horses, cats and mice (Figure 1A and supplementary Figure S1). Interestingly, substitution with tyrosine (Y) in this position appears specifically in *Catarrhini* group, including the old-world monkeys and apes (Figure 1B).

**Figure 2.**
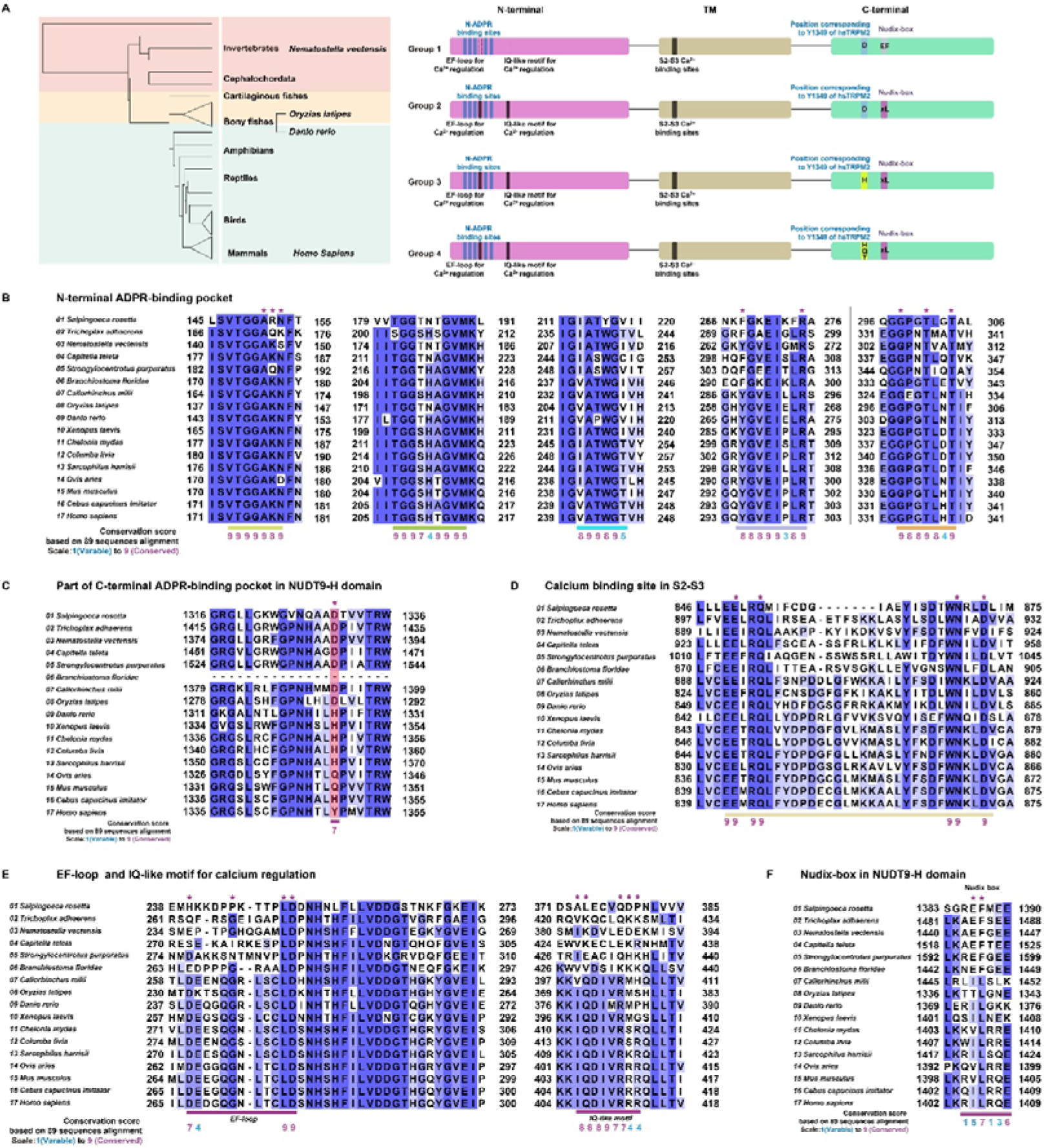
Summary and sequence alignments showing the motifs involved in activation of 17 representative TRPM2 channels by ADPR and calcium. (A) Summary of changes in the motifs involved in TRPM2 channel activation by ADPR and calcium among different groups and during evolution. (B-F) Sequence alignments showing the motifs involved in channel activation in 17 representative TRPM2 sequences by ADPR and calcium.

**Figure 3.**
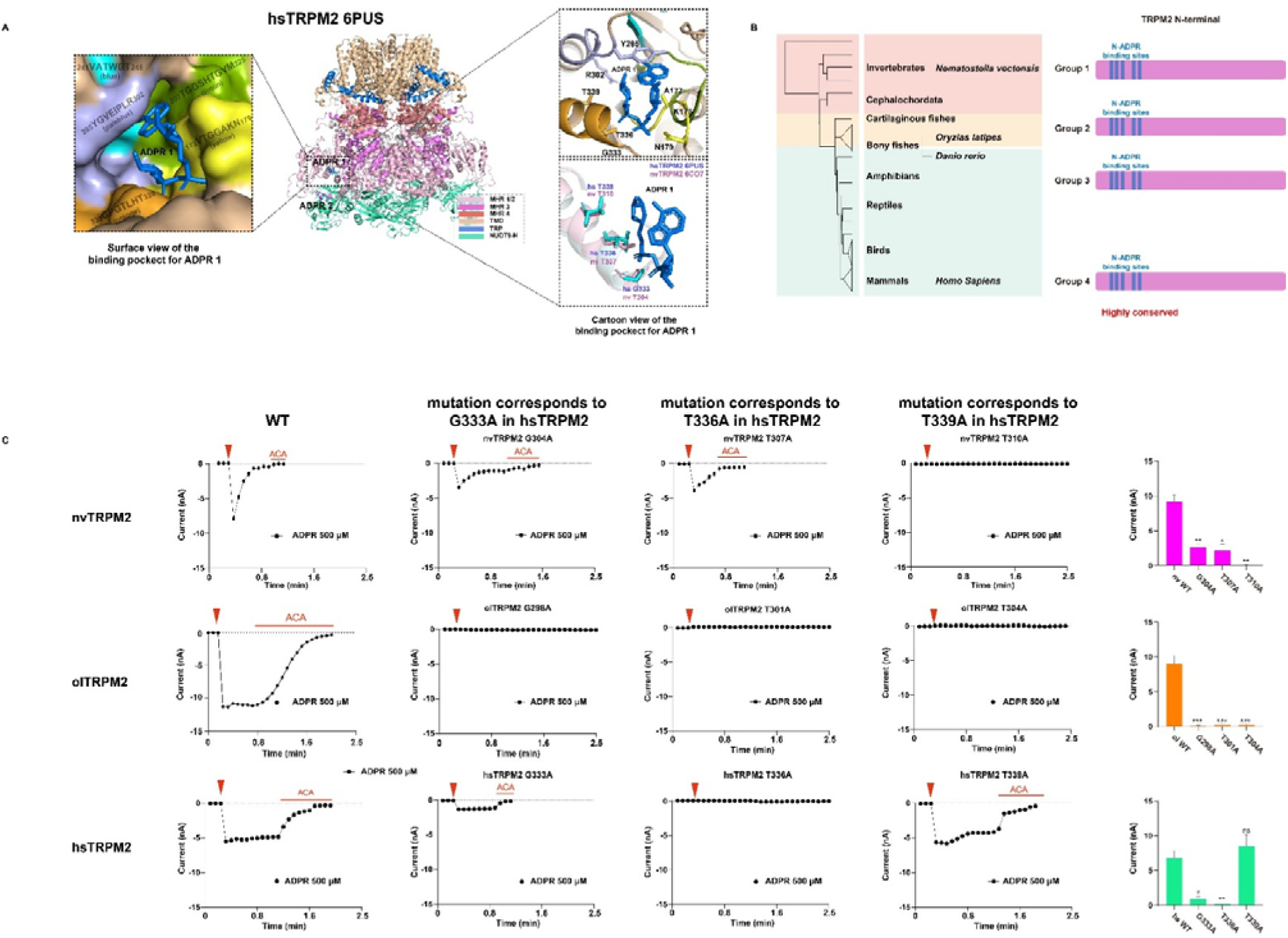
The N-terminal ADPR-binding pocket is critical for ADPR-induced activation of TRPM2 channels from invertebrates to higher animals. (A) Cartoon and surface view of the location and residues of the N-terminal ADPR-binding pocket in hsTRPM2 channel (6PUS). (B) Summary of the residues forming or around the N-terminal ADPR-binding pocket of TRPM2 channels evolved among different groups and the corresponding mutagenesis studies. (C) Representative whole-cell patch-clamp recordings showing inward currents at -80 mV induced by 500 μM ADPR from HEK293T cells expressing WT or indicated mutant nvTRPM2, olTRPM2 or hsTRPM2 channel. The red triangle in each panel indicates the time point at which whole-cell configuration was established. Voltage ramps from -100 mV to 100 mV were applied every 5 s. The red bars represent application of 20 μM N-(p-amylcinnamoyl) anthranilic acid. Summary of mean ± SEM current densities recorded from the number of cells in each case (n = 3-8) is shown on the far right.

The ADPR catalytic ability of the NUDT9-H domain in TRPM2 have been reported in some species [24, 25]. Previous studies also suggest the change of EF residues to IL in the Nudix box within this domain causes loss of the enzymatic activity in some species including hsTRPM2 (_1405_IL_1406_) [34, 40]. We found that these two residues were variable among species (Supplementary Figure S7). The EF motif is present in invertebrate TRPM2, but replaced with xL, with IL in most vertebrate species since the emergence of cartilaginous fishes. This event happened much earlier (Figure 1B, 2A, 2F and supplementary Figure S2) than the initial appearance of H residue in the position corresponding to Y1349 in hsTRPM2.

### 3.3 Evolution of TRPM2 on sequence level: multiple motifs for Ca^2+^ regulation of TRPM2 were acquired in vertebrates

Intracellular Ca^2+^ is critical in hsTRPM2 channel activation [28, 29], and Ca^2+^ binding is likely coordinated by three separate Ca^2+^-binding sites, the Ca^2+^-binding site formed by the S2-S3 helix, the Ca^2+^/CaM-binding IQ-like motif and the EF-loop [30–33]. We further investigated the conservation patterns of these Ca^2+^-binding sites in all species.

Based on the atomic structure of nvTRPM2 (6CO7) solved in the presence of Ca^2+^ [30], Ca^2+^ interacts with four residues, namely, E893, Q896, N918 and D921 in the S2-S3 helix of nvTRPM2 (Figure 2A, 2D). These four residues are also present in hsTRPM2 and, in fact, completely conserved in all 89 sequences analyzed (supplementary Figure S8), indicating that this Ca^2+^-binding site persists through billion years.

The IQ-like motif (IQxxxRRRxxL) is present in TRPM2 from vertebrates (Figure 2A, 2E and Supplementary Figure S9), but absent in TRPM2 of invertebrates and chordate such as *Branchiostoma floridae* (Figure 1B and Figure 2A). Therefore, unlike the TRPM2 channels in vertebrates, the primitive TRPM2 channels in invertebrates does not need this Ca^2+^/CaM-binding motif as a mechanism for channel activation by Ca^2+^.

Unlike the highly conserved Ca^2+^-binding site in the S2-S3 helix, the EF-loop (DExxxGxxxxLD) is absent in TRPM2 channels from invertebrates and ancient vertebrates. This motif was acquired in vertebrates around 460-500 million years ago (Figure 1B and supplementary Figure S2) and retains in almost all vertebrates (Figure 2A, 2E and Supplementary Figure S1, S9). Of note, the last two residues, LD, in the EF-loop previously reported to be involved in the activation of hsTRPM2 channel by Ca^2+^ [33], are highly conserved in invertebrates, albeit loss of the EF-loop sequence.

### 3.4 Evolution of TRPM2 on sequence level: three identifiable stages of evolutionary changes in TRPM2 domains involved in activation by ADPR and _Ca_2+

Based on the conservation patterns of the functional important motifs and domains, we propose three identifiable evolutionary stages in terms of TRPM2 channel activation by ADPR and Ca^2+^ (Figure 1A, 2B). The first stage happened during the transition from invertebrates to vertebrates (after the emergence of *Branchiostoma floridae,* a chordate), with change of the catalytic EF residues in the Nudix box from EF to xL and acquirement of the EF-loop and IQ-like motifs. The second stage occurred during the water-to-land transition in vertebrates about 350-430 million years ago, during which the position corresponding to Y1349 in the hsTRPM2 changed from D in invertebrates and fish to H in tetropods. The third stage happened roughly 160 million years ago, during which Y and Q emerged at the position corresponding to Y1349 in the hsTRPM2.

According to these evolutionary stages, we can divide all TRPM2 into four groups (Figure 2A). Of note, the groups mentioned here are based on whether they possess the sequence residues discussed above (Figure 2A). TRPM2 channels possessing the corresponding “signatures” sequence patterns/residues are classified as one group. The first group existed before the 1^st^ stage and includes TRPM2 from all invertebrates and *Branchiostoma floridae*, with the nvTRPM2 channel as an example. The second group consists of all TRPM2 channels between the 1^st^ and 2^nd^ stages, encompassing most fish species, with the *Oryzias latipes* TRPM2 (olTRPM2) as a representative. The third group is composed of TRPM2 channels from amphibians, reptiles, birds and some mammalian. This group has H at the position corresponding to Y1349 in the hsTRPM2, and one example of this group is drTRPM2 (*Danio rerio*, zebrafish TRPM2). The 4^th^ group contains TRPM2 channels emerging after the 3^rd^ stage, with Q or Y at the position corresponding to Y1349 in the hsTRPM2, with the mmTRPM2 (*Mus musculus*, mouse TRPM2) and hsTRPM2 as representatives. Next, we experimentally examined channel activations by ADPR and Ca^2+^ in the representative TRPM2 channels in each of the four groups.

### 3.5 Functional characterizations: the N-terminal ADPR-binding pocket is critical for ADPR-induced activation of TRPM2 channels from invertebrates to vertebrates

The sequence conservation usually indicates function importance. To examine whether the highly conserved N-terminal ADPR-binding pocket (Figure 2B) is indeed functionally indispensable, especially for the previously uncharacterized olTRPM2, we performed the following investigations.

All TRPM2 channels that have been characterized so far can be activated by ADPR, albeit with some differences in the reported EC_50_ value (50% effective concentration), 2.3 ± 0.2 μM for the nvTRPM2 channel [41] and 40.0 ± 4.5 μM for the hsTRPM2 channel [42]. In this study, we showed the EC_50_ value of 1.1 ± 0.8 μM for the olTRPM2 channel (supplementary Figure 3E), which is close to that for the nvTRPM2 channel and ∼40 times lower than that for the hsTRPM2 channel. The residues forming the N-terminal ADPR-binding pocket (ADPR1) in the hsTRPM2 channel structure, are highly conserved (Figure 2B), supporting its functional importance in TRPM2 channel activation by ADPR. Consistently, alanine replacement of some of these conserved residues, including R302 and R358 in the hsTRPM2 channel and R278 and R358 in the drTRPM2 channel, resulted in loss of channel activation by ADPR (supplementary Table S3).

To further assess the functional importance of three conserved residues near the N-terminal ADPR-binding pocket corresponding to G333, T336 and T339 in the hsTRPM2 channel (Figure 3A, 3B and supplementary Table S3), we introduced alanine substitution into these positions. Our results showed that all the mutations impaired channel activation by ADPR. There was complete loss of activation in response to application of 500 μM ADPR, a saturating concentration for the WT channel, as a result of T336A in the hsTRPM2, G298A, T301A and T304A in the olTRPM2, and T310A in the nvTRPM2 (Figure 3C). In the hsTRPM2, while the T339A mutation had no effect, the G333A mutation strongly reduced ADPR-induced currents (Figure 3C). In the nvTRPM2 channel, both the G304A and T307A mutations led to decreased ADPR-induced currents. Among the mutants showing no current response to 500 μM ADPR (Figure 3C), they also failed to respond to 5 mM ADPR (supplementary Figure S3A-D), with an exception of the olTRPM2 T301A mutant channel that mediated a very small current, albeit no significant reduction in their cell surface expression (supplementary Figure S5B, S5C and S5E). These results, while revealing some differences in the contribution of individual residues, support the critical role of this N-terminal ADPR-binding pocket in mediating ADPR activation of TRPM2 channels from early metazoan (nvTRPM2) and vertebrates (olTRPM2 and hsTRPM2).

### 3.6 Functional characterizations: the mutation of residue corresponding to Y1349 in hsTRPM2 indicates the necessity of the NUDT9-H domain for TRPM2 activation by ADPR

As shown in our previous study [42], Y1349 in the C-terminus of the hsTRPM2 (Figure 4A) plays an important role in channel activation by ADPR (supplementary Table S3). To investigate the effect of the amino acid substitutions that occurred in evolutionary history on the channel activation by ADPR, we examined mutant hsTRPM2 channel with Y replaced with D, H or Q that is present in the equivalent position of the nv, ol, dr and mm TRPM2 channels, respectively (Figure 4B). All these mutants showed similar cell surface expression to the WT hsTRPM2 (supplementary Figure S5E). Interestingly, the Y1349D mutant was only weakly activated by ADPR (Figure 4C), with 5 mM evoking considerably reduced currents (supplementary Figure S3A-D), suggesting a deleterious effect for reversal mutation of Y1349 to its ancestral state (i.e. D) [43]. This was also true for the H residue.

**Figure 4.**
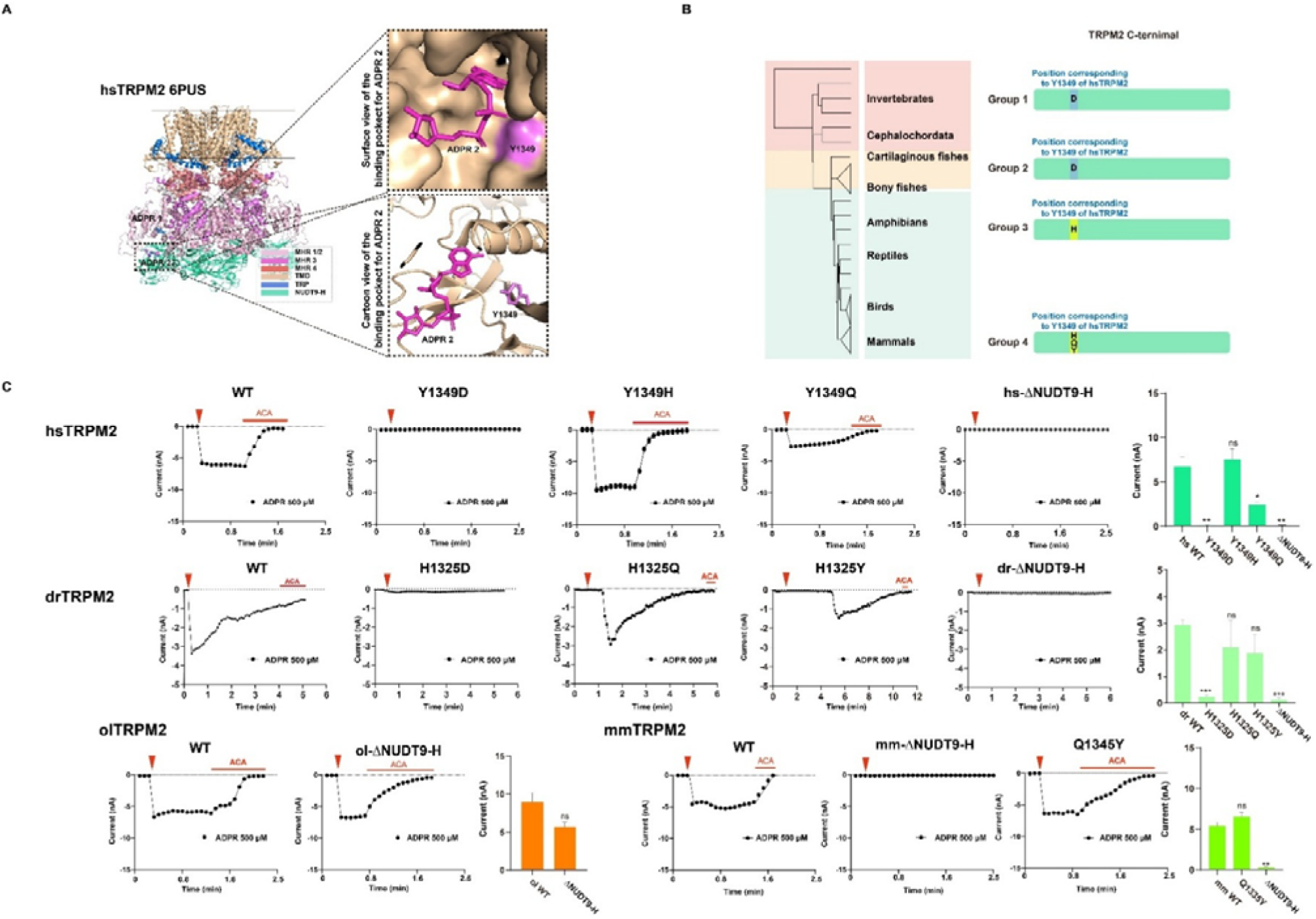
The residue corresponding to Y1349 in hsTRPM2 indicates the necessity of the NUDT9-H domain for TRPM2 channel activation by ADPR. (A) Cartoon and surface view of hsTRPM2 (6PUS) showing the location and residues of the C-terminal ADPR-binding pocket and residue Y1349. (B) Summary of the residues forming or around the C-termini ADPR binding pocket of TRPM2 channels evolved among different groups and the corresponding mutagenesis studies. (C) Representative whole-cell patch recordings showing inward currents at -80 mV induced by 500 μM ADPR from HEK293T cells expressing WT or indicated mutant hsTRPM2, drTRPM2, olTRPM2 or mmTRPM2 channel. The red triangle in each panel indicates the time point at which whole-cell configuration was established. Voltage ramps from -100 mV to 100 mV were applied every 5 s. The red bars represent application of 20 μM N-(p-amylcinnamoyl) anthranilic acid. Summary of mean ± SEM current densities recorded from the number of cells in each case (n = 3-8) is shown on the right.

H1325 in the drTRPM2 corresponds to Y1349 in the hsTRPM2, based on the sequence alignments and structural comparisons. Introduction of the reversal mutation D (i.e. H1325D) in the drTRPM2 channel also significantly reduced ADPR-induced currents (Figure 4C), while mutations to the “tetropod” form, i.e. H1325Q and H1325Y in the drTRPM2 channel barely affected ADPR-induced activation (Figure 4C). All the drTRPM2 mutants exhibited a surface expression level comparable to the WT drTRPM2 (supplementary Figure S5D). These results suggest that any loss of or impact on TRPM2 channel activation is not attributable to variations in surface expression levels. Moreover, concerning the H1325Y mutant drTRPM2, even though its surface expression level and current magnitude were comparable to that of the WT drTRPM2 (supplementary Figure S5D; Figure 4C), its activation was notably slower. We conducted additional experiments using 5 mM ADPR (supplementary Figure S3E), and observed that the activation of the H1325Y mutant was faster than that evoked by 500 μM ADPR, but was still slower than that of the WT and other mutants.

The residue corresponding to Y1349 in the hsTRPM2 channel is the only residue in this region showing a clear conservation pattern. Our results suggest that the nvTRPM2 channel has D at this position (Figure 4B) and shows no activation by ADPR, which is consistent with the observation that the activation of the nvTRPM2 channel by ADPR does not require the C-terminal NUDT9-H domain (supplementary Table S3) [41]. We hypothesized that the C-terminal NUDT9-H domain is not necessary for ADPR-induced activation of a TRPM2 channel having D at this position. Indeed, deletion of the NUDT9-H in the olTRPM2 channel, which also has D at this position like TRPM2 in most of fish (Figure 3B), had no effect on channel activation by ADPR (Figure 4C). On the other hand, introduction of the Y1349H mutation in the hsTRPM2 channel barely altered channel activation by ADPR (Figure 4C), suggesting the NUDT9-H domain carrying H or Y is required for mediating ADPR binding and channel activation. This notion is consistent with the crucial role of the NUDT9-H domain in the activation by ADPR of the drTRPM2 channel having H, and the hsTRPM2 channel bearing Y (Figure 4B and supplementary Table S3) [25, 44, 45]. Furthermore, introduction of the Y1349Q mutation led to reduced activation of the hsTRPM2 channel by ADPR (Figure 4C and supplementary Figure S3A). The mmTRPM2 contains Q at the corresponding position and replacement of Q with Y increased ADPR-induced currents (Figure 3C). Therefore, we propose that the NUDT9-H domain carrying Q at this position is equally critical for TRPM2 channel activation by ADPR. In supporting this notion, deletion of the NUDT9-H domain in the mmTRPM2 channel resulted in loss of activation by ADPR (Figure 4C, supplementary Figure S3A and supplementary Table S3). Therefore, for TRPM2 channels having a D at the position corresponding to Y1349 in the hsTRPM2, their NUDT9-H domain is not necessary for channel activation by ADPR (Figure 4B and supplementary Table S3). However, the NUDT9-H domain is indispensable for the activation by ADPR of a TRPM2 channel having H, Q or Y at such a position (Figure 4B and supplementary Table S3).

### 3.7 Functional characterizations: the conserved Ca^2+^-binding site at the S2-S3 helix is indispensable for Ca^2+^-induced activation of all TRPM2 channels

Intracellular Ca^2+^ can activate the hsTRPM2 channel independently of or in synergy with ADPR [28, 29]. The Ca^2+^-binding site formed by the S2-S3 helix (Figure 5A) [25, 30] are highly conserved (Figure 5B, supplementary Figure S1, S8, and supplementary Table. S4), indicating its critical role in channel activation by Ca^2+^, as previously shown in the nvTRPM2, drTRPM2 and hsTRPM2 channels [25]. Here, to determine whether this highly conserved Ca^2+^-binding site also plays a critical role especially for the previously uncharacterized TRPM2 channels, we examined its role in Ca^2+^-induced activation of the olTRPM2 channel. Mutation of E828, Q854 or Q857 to alanine resulted in no channel activation by ADPR in the absence of Ca^2+^ (Figure 5C), and reduced activation in the presence of Ca^2+^ (supplementary Figure S4A). These results support the notion that this highly conserved Ca^2+^-binding site is functionally indispensable in both primitive and sophisticated TRPM2 channels.

**Figure 5.**
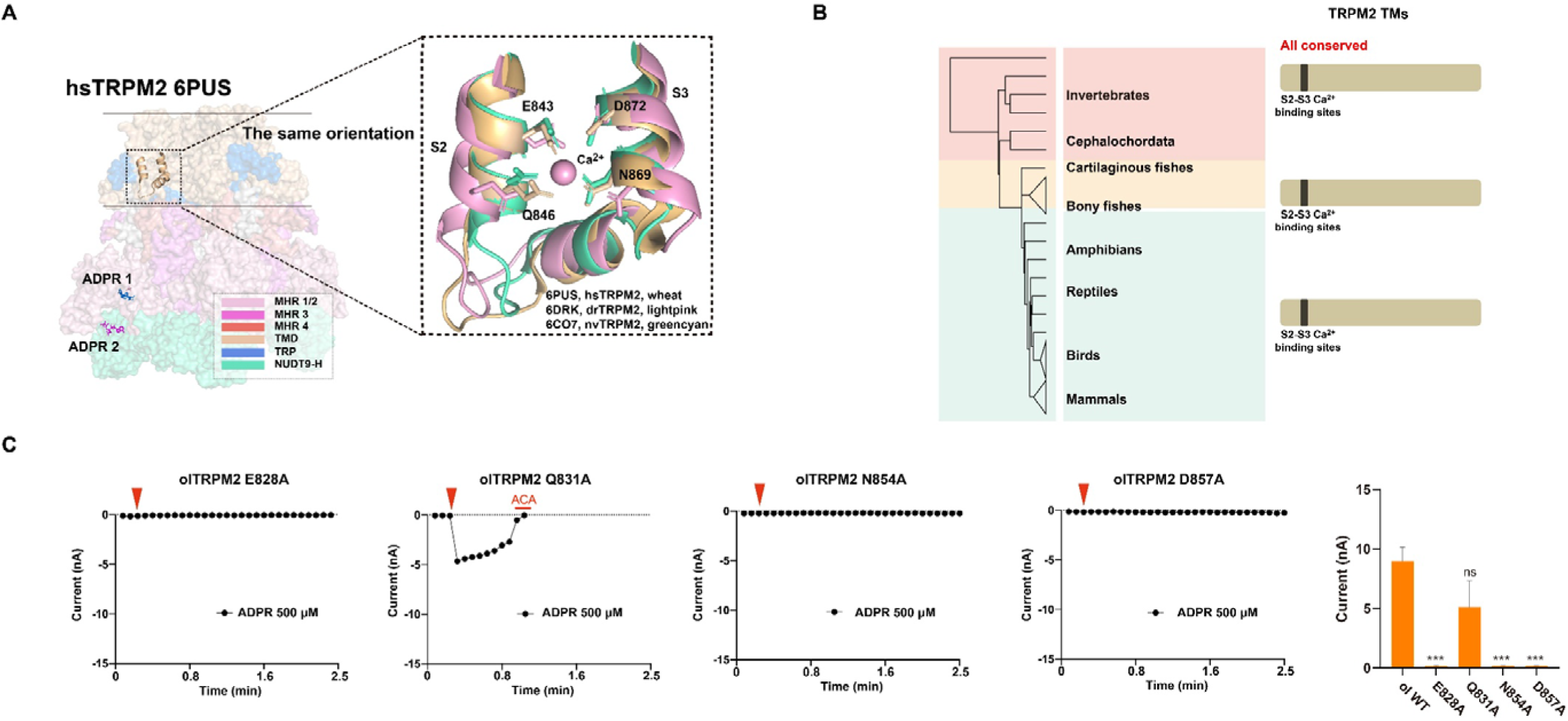
Ca^2+^-binding site in the S2-S3 helix is indispensable for Ca^2+^-induced TRPM2 channel activation. (A) Cartoon of hsTRPM2 (6PUS, wheat), drTRPM2 (6DRK, light pink) and nvTRPM2 (6CO7, greencyan) showing the location and residues involved in Ca^2+^ binding. (B) Sequence alignments of the S2-S3 amino acid sequences showing residues involved in Ca^2+^ binding to TRPM2 channels from 17 representative species. (C) Summary of the S2-S3 residues involved in Ca^2+^ binding to TRPM2 channels evolved among different groups and the corresponding mutagenesis studies. (D) Representative whole-cell patch-clamp recordings showing inward currents at -80 mV induced by 500 μM ADPR from HEK293T cells expressing indicated mutant olTRPM2 channel. The red triangle in each panel indicates the time point at which whole-cell configuration was established. Voltage ramps from -100 mV to 100 mV were applied every 5 s. The red bars represent application of 20 μM N-(p-amylcinnamoyl) anthranilic acid. Summary of mean ± SEM current densities recorded from the number of cells in each case (n = 3-8) is shown on the right.

### 3.8 Functional characterizations: the evolution of TRPM2 in channel regulation and activation by Ca^2+^

The EF-loop and IQ-like motif in the N-terminus, as introduced above, known to participate in the regulation and activation of the TRPM2 channel by Ca^2+^, are absent in invertebrate TRPM2, but such sequence patterns acquired before the emergence of *Callorhinchus milii* (Figure 2E, 6B and 6D). Existing evidence shows the cooperation of intracellular ADPR and Ca^2+^ in channel activation is different for the nvTRPM2 and hsTRPM2 channels [42]. Removal of intracellular basal Ca^2+^ using EGTA barely affected the nvTRPM2 channel activation, but significantly reduced the hsTRPM2 channel activation by ADPR (Figure 6C). We further investigated the two fish TRPM2 representatives. Surprisingly, the olTRPM2 channel behaved like the nvTRPM2 channel in that intracellular Ca^2+^ is not necessary for channel activation by ADPR (Figure 6C). In contrast, the drTRPM2 channel behaved like the hsTRPM2 channel, for its activation strongly depends on the cooperation of ADPR and Ca^2+^ (Figure 6C). These observations indicate that, the primitive or ancient TRPM2 channels such as nvTRPM2 may not require intracellular Ca^2+^ for ADPR-induced activation, while the sophisticated TRPM2 channels such as hsTRPM2 strongly depend on the cooperation of intracellular ADPR and Ca^2+^ for channel activation. This event should happen before the divergence of bony fish.

**Figure 6.**
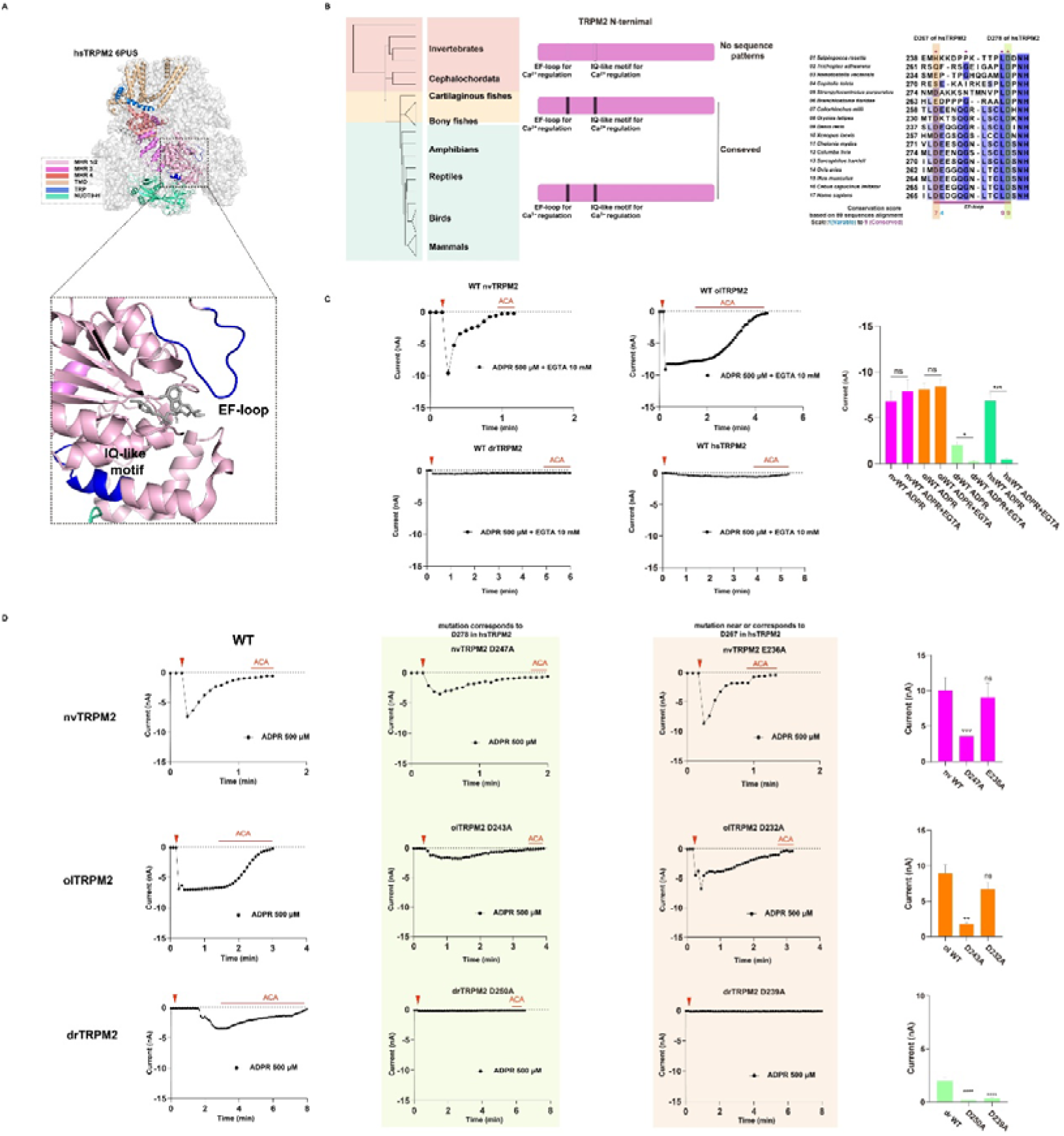
Different sequence pattern other than EF-loop used in regulating calcium-induced activation of nvTRPM2 and other ancient TRPM2s. (A) Cartoon and surface view of hsTRPM2 channel (6PUS) showing the location of EF-loop and IQ-like motif which contribute to calcium regulation of the channel. (B) Summary of residues of, or corresponding to, EF-loop and IQ-like motif in TRPM2 channels evolved among different groups and the corresponding mutagenesis studies. (C) Representative whole-cell patch recordings showing inward currents at -80 mV induced by 500 μM ADPR with 10 mM ethylene glycol tetraacetic acid (EGTA) inside the cell from HEK293T cells expressing WT nvTRPM2, olTRPM2, drTRPM2 or hsTRPM2 channel. (D) Representative whole-cell patch recordings of inward currents at -80 mV induced by 500 μM ADPR with additional 1 mM Ca^2+^ from HEK293T cells expressing WT or indicated mutant nvTRPM2, olTRPM2 or drTRPM2 channel. (C) (D) The red triangle in each panel indicates the time point at which whole-cell configuration was established. Voltage ramps from -100 mV to 100 mV were applied every 5 s. The red bars represent application of 20 μM N-(p-amylcinnamoyl) anthranilic acid. Summary of mean ± SEM current densities recorded from the number of cells in each case (n = 3-8) is shown on the right.

The residue corresponding to D278 in the EF-loop of the hsTRPM2 channel is highly conserved across all species (Figure 6B and 6D). Introduction of alanine substitution in the hsTRPM2 channel led to channel inactivation [33]. Alanine mutation of the equivalent D243 in the olTRPM2 channel and D250 in the drTRPM2 channel led to reduced sensitivity to Ca^2+^ (Figure 6D), suggesting that this residue in the olTRPM2 and drTRPM2 channels is also critical for Ca^2+^-induced channel activation. Similarly, introduction of the D247A mutation in the nvTRPM2 channel reduced channel activation (Figure 6D). These results consistently support the notion that even though the sequence pattern of the EF-loop is absent in the primitive TRPM2 channels such as nvTRPM2 (Figure 2B), this domain, particularly the conserved D residue, has an important role in Ca^2+^-induced channel activation.

The negatively-charged residues in the EF-loop are usually involved in Ca^2+^ binding. Therefore, we investigated such residues corresponding to D267 in the hsTRPM2 channel, although less conserved (Figure 6D). Introduction of D267A mutation in the hsTRPM2 channel, despite without abolishing channel activation by Ca^2+^, reduced channel activation (Figure 6B and supplementary Table S3). The drTRPM2 channel behaved like the hsTRPM2 channel, with significantly reduced activation of the D239A mutant channel (Figure 6D), while the membrane surface expression levels weren’t altered (supplementary Figure S5D). Alanine substitution of the corresponding D232 in the olTRPM2 channel barely affected channel activation (Figure 6D supplementary Table S3). Invertebrate TRPM2 channels such as nvTRPM2 did not possess the EF-loop sequence pattern. In the nvTRPM2 channel, there is a negatively-charged residue E236 near the residue corresponding to D267 in the hsTRPM2 channel but introduction of alanine mutation did not affect channel activation in the presence of Ca^2+^ (Figure 6D and supplementary Table S3). Therefore, D267 in the hsTRPM2 channel and D239 in the drTRPM2 channel, despite being less conserved, are important for channel activation by Ca^2+^, whereas the negatively-charged residues corresponding to or near D232 in the olTRPM2 or E236 in the nvTRPM2 channel is of less importance. These results also suggest that the contributions of certain residues or motifs are different for TRPM2 channels from different species.

## 4. Discussion and Conclusion

In this study, we systematically analyzed TRPM2 channels from more than 280 species focusing on the evolutionary trajectory of channel activation modes by ADPR and Ca^2+^. By combining molecular evolution approaches with mutagenesis and patch-clamp recording, we provide both analytical and experimental evidences to show the following. First of all, the N-terminal ADPR-binding pocket was present since the emergence of TRPM2 channels and it is highly conserved and critical for ADPR-induced activation of the channels from invertebrates to vertebrates. Secondly, the C-terminal NUDT9-H domain, though also present in ancient TRPM2 channels, is not required for channel activation by ADPR until roughly 350-430 million years ago but since then, became indispensable for channel activation by ADPR. We characterized the olTRPM2 channel, as an intermediate from the ancient TRPM2 channel like the nvTRPM2 to the TRPM2 channels that evolved more recently such as the drTRPM2, mmTRPM2 and hsTRPM2 channels. For the olTRPM2 channel, the NUDT9-H domain is not essential for ADPR activation, similar to what was previously reported for the nvTRPM2 channel [24], and in stark contrast with its close relative drTRPM2 (Figure 4B, 4C and supplementary Table S3). Thirdly, for the olTRPM2 channel, intracellular Ca^2+^ is not necessary for its activation by ADPR, which behaves like the nvTRPM2 channel. In contrast, activation of its close relative drTRPM2 and hsTRPM2 channels requires the cooperation of intracellular ADPR and Ca^2+^ (Figure 6C). Fourthly, among the three identifiable Ca^2+^-binding domains, the binding site in the S2-S3 helix is highly conserved and present since the beginning, which is crucial for channel activation by Ca^2+^. The IQ-like motif and the EF-loop, which are involved in the activation of TRPM2 channels, are less conserved and present after the emergence of vertebrates; the primitive TRPM2 channels in invertebrates did not need these two Ca^2+^-binding sites for channel activation by Ca^2+^; for the sophisticated TRPM2s, mutational disruption of any one of the three Ca^2+^-binding sites affected the activation of TRPM2 channels (Table S3), indicating that the mutation or loss of any of these domains impacts channel activation by Ca^2+^. Finally, we have identified important permissive changes at the position corresponding to Y1349 in the hsTRPM2 channel that occurred during the transition from water to land transition in vertebrates. The occurrence of H, Q or Y at this position indicates the origin of the dependence of the C-terminal NUDT9-H domain for ADPR-induced channel activation in tetropods. In summary, our study reveals that the species differences in the mode or structural requirements of TRPM2 channel activation by ADPR and Ca^2+^ reflect evolutionary changes in the binding domain and motifs for ADPR and Ca^2+^.

Here, we characterized the evolution in the TRPM2 channel activation mode by ADPR and Ca^2+^, which happened in at least three identifiable stages (Figure 7A). Based on the activation mode, all TRPM2 channels can be classified as “primitive”, “intermediate” and “sophisticated” (Figure 7B). The activation model for the primitive TRPM2 channels, represented by the nvTRPM2, is composed of an ADPR-binding site in the N-terminal MHR1/2 domain and a Ca^2+^-binding site in the S2-S3 helix. These TRPM2 channels are present in invertabrates and *Branchiostoma floridae*, before the 1^st^ evolutionary stage (Figure 7A). The NUDT9-H domain in the TRPM2 channels shares approximately 40% sequence identity to NUDT9 [27], a pyrophosphatase in the mitochondria that catalyzes hydrolysis of ADPR to AMP and ribose-5-phosphate [40]. The TRPM2 channels in invertebrate species choanoflagellate (*Salpingoeca rosetta*) and sea anemones (*Nematostella vectensis*) retain the pyrophosphate activity, and function as a chanzyme [24, 25]. This catalytic activity of the NUDT9-H domain was lost in the hsTRPM2 channel, likely due to replacement of the EF residues with IL in the canonical Nudix-box motif [34]. Here, in the primitive TRPM2 channels, the C-terminal NUDT9-H domain possesses the EF residues in the Nudix box (Figure 2F) and exhibits a catalytic activity, but this domain is not necessary for channel activation by ADPR, and there is lack of the EF-loop and IQ-like motif for channel activation by Ca^2+^ (Figure 2E and Supplementary Figure S1). It could be activated by ADPR without the cooperation of intracellular Ca^2+^ (Figure 6C). Therefore, the activation mode of the primitive TRPM2 channels by ADPR and Ca^2+^ can be described as “1+1+C”, with C referring to a chanzyme. A recent study investigating the cryo-EM structures and the activation of the most ancient TRPM2, *Salpingoeca rosetta* (sr) TRPM2) suggests that the channel lacking the NUDT9-H domain is activated by ADPR [46], which is highly consistent with our findings.

**Figure 7.**
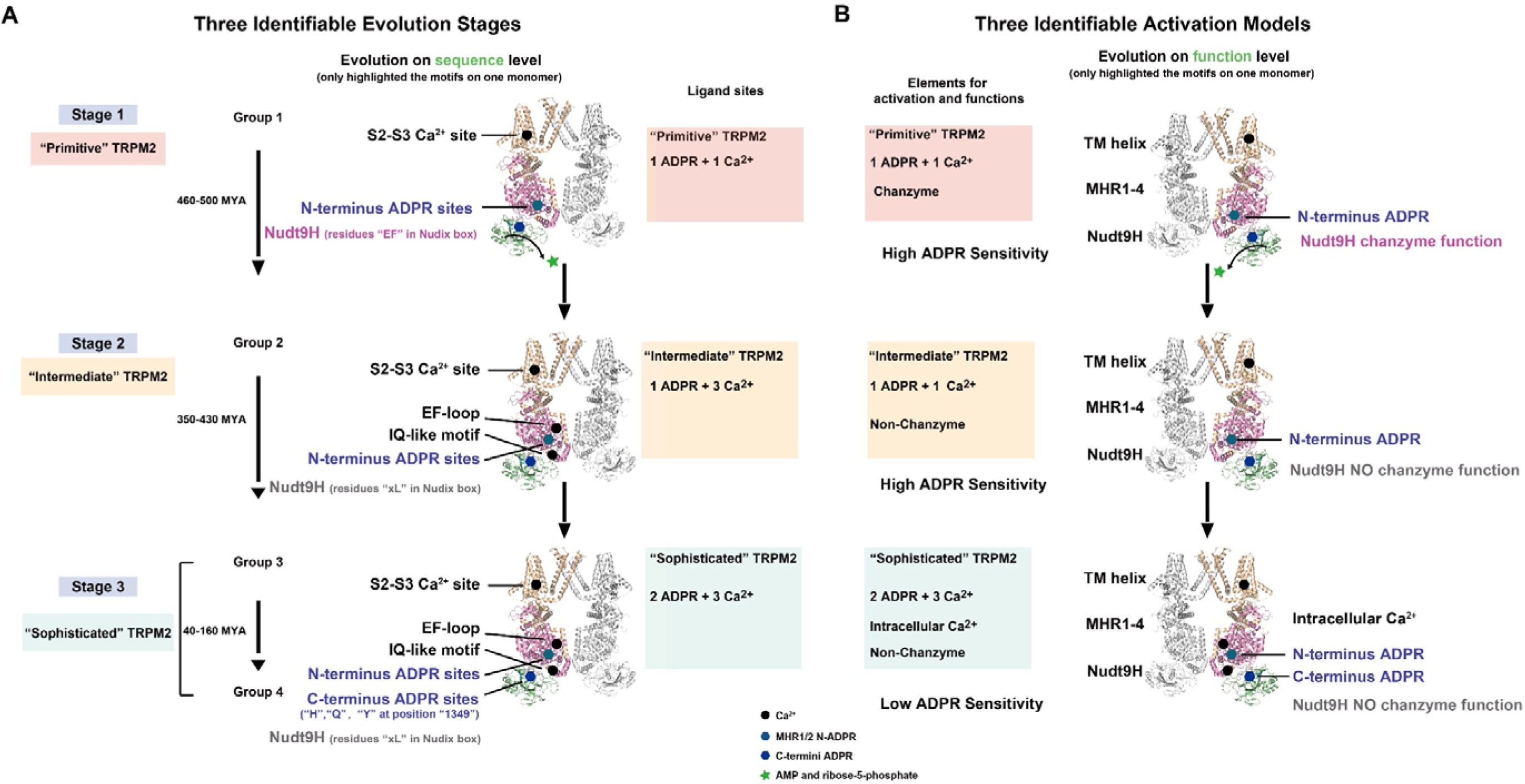
The TRPM2 channel activation models and evolution stages. The designation of activation modes and evolution stages (A) and the details of each activation mode (B).

The nvTRPM2 channel and its mutant without the NUDT9-H domain displayed a higher sensitivity to ADPR than the hsTRPM2 channel [24, 41, 47]. The pyrophosphatase activity of the NUDT9-H domain in the nvTRPM2 and other ancient TRPM2 channels [34] is likely high [24, 34] and the NUDT9-H domain may serve a mechanism to degrade ADPR. In the primitive TRPM2 channels, ADPR has dual functions, one as a ligand for channel activation and the other as a substrate for NUDT9-H-mediated hydrolysis. *Therefore, the primitive TRPM2 channels, albeit showing a higher ADPR sensitivity, have a low efficiency using ADPR as an activation ligand or agonist.* ROS such as H_2_O_2_ activates the TRPM2 channel indirectly through promoting the production of ADPR [48]. Interestingly, for the nvTRPM2 channel, H_2_O_2_ was less able to activate the WT channel than the mutant channel with the NUDT9-H domain [47, 49, 50], supporting the notion that degradation of ADPR slows down the accumulation of ADPR for channel activation.

The intermediate TRPM2 channels emerged after the 1^st^ evolutionary stage that are present in most of fishes that evolved roughly 460-500 million years ago (Figure 1B, Figure 7 and supplementary Figure S2). At the sequence level, compared to the primitive TRPM2 channels, the intermediate TRPM2 channels have acquired two additional motifs for Ca^2+^ binding (i.e. EF-loop and IQ-like motif; Figure 7A). At the biophysical function level, the contribution of EF-loop, particularly the first negatively charged D residue, is not required for channel activation (Figure 6D). Furthermore, activation of the intermediate TRPM2 channels by ADPR does not require intracellular Ca^2+^ (Figure 6C). Therefore, the channel activation mode can be summarized as one ADPR and one Ca^2+^-binding site with loss of the pyrophosphatase enzyme activity, or “1+1+0” model, as shown in this study for the olTRPM2 channel. Channel activation depends on ADPR binding to the N-terminal MHR1/2 domain and Ca^2+^ binding to the S2-S3 helix. The NUDT9-H domain of the intermediate TRPM2 channels loses its pyrophosphatase activity, due to substitution of the EF residues with xL in the Nudix box [24, 25, 34] (Figure 2F and supplementary Figure S7B), and it is not required for channel activation. The advantage arising from loss of the ADPR-degrading ability is that the intermediate TRPM2 channels use ADPR more efficiently as a ligand, with retaining the ADPR sensitivity, as suggested by the observation that the olTRPM2 channel has an ADPR EC_50_ (supplementary Figure S3F) that is similar to that at the nvTRPM2 channel [24, 41]. In addition, deletion of the NUDT9-H domain in the olTRPM2 channel did not affect channel activation by ADPR, indicating that the NUDT9-H domain adopts an insensitive form, probably related with the D at the position corresponding to Y1349 in the hsTRPM2 channel (Figure 4C). During the evolution, the intermediate TRPM2 channels accumulated other changes, including the EF-loop and IQ-like motif (Figure 6B and Supplementary Figure S1), leading to gain of an additional level of regulation or activation by Ca^2+^.

The sophisticated TRPM2 channels appeared after the 2^nd^ evolutionary stage roughly 350-430 million years ago (Figure 1B, Figure 7 and supplementary Figure S2). They contain all the TRPM2 channels from the third and fourth groups, including the drTRPM2 channel and TRPM2 channels from amphibians, reptiles, birds and mammals. Their activation is coordinated by two ADPR-binding sites and three Ca^2+^-binding sites, but no chanzyme function, or the “2+3+0” model. On the biophysical function aspects, their activation requires the cooperation of intracellular ADPR and Ca^2+^ (Figure 6C). One of the salient changes is that the NUDT9-H domain becomes indispensable for channel activation by ADPR (Figure 4 and supplementary Table S3). Loss of the N-terminal ADPR-binding site or the C-terminal NUDT9-H domain results in no channel activation by ADPR, as shown in the drTRPM2, hsTRPM2 [25, 44, 45] and mmTRPM2 channels (Figure 4B, 4C and supplementary Table S3). The sophisticated TRPM2 channels also require the Ca^2+^-binding sites in the S2-S3 helix, the EF-loop and IQ-like motif for channel activation by Ca^2+^, which is similar to the intermediate TRPM2 channels. Therefore, the sophisticated TRPM2 channels have the most complicated activation mode and depend on the cooperation of intracellular ADPR and Ca^2+^ (Figure 7B).

As demonstrated in this study, a key change in the NUDT9-H domain of the sophisticated TRPM2 channels is that they acquire the H residue at the position corresponding to 1349 in the hsTRPM2 channel that replaces D in the primitive TRPM2 channels. H emerged first at 350∼430 million years ago (Figure 1B and supplementary Figure S2), and further replaced by Q or Y at about 160 million years ago (Figure 1B and supplementary Figure S2). We demonstrated that Y1349 was a permissive mutation, making reversion to ancestral state (i.e. D) deleterious (Figure 4C) [43]. H and Q at this position are similar to Y, being permissive [43] and allowing the acquisition of mutations that cause partial function loss of the N-terminal ADPR-binding site, and the C-terminal NUDT9-H domain becomes indispensable for channel activation by ADPR.

The conserved N-terminal ADPR-binding site plays an important role in channel activation by ADPR since the emergence of the first TRPM2 channel, as substitution of one residue in this site in the nvTRPM2, olTRPM2, drTRPM2 and hsTRPM2 channels abolished channel activation by ADPR (Figure 3B) [25]. The requirement of the C-terminal NUDT9-H domain for channel activation by ADPR occurred more than 600 million years later, upon the emergence of H at the position corresponding to Y1349 in the hsTRPM2 channel. Loss of its pyrophosphatase activity, occurring roughly 100 million years earlier, did not confer the necessity of the NUDT9-H domain for channel activation by ADPR (Figure 1B, 7B).

It is well-known that the accumulation of O_2_ in the environments affected largely the trajectory of evolution, facilitated adaptations within oxygen-sensing and intrinsic regulatory pathways [51, 52]. Studies of male Wistar rats exposed to different levels of oxygen suggest that exposure to high oxygen concentrations can induce excessive levels of oxidative stress in rats [53]. Based on the climate changes during these 1000 million years (Figure 1B), the oxygen levels may serve as a selective pressure during the first two stages in the evolution of the TRPM2 channel activation mode, and further studies are required to validate such a possibility.

In conclusion, we analyzed the evolutionary changes in the activation mode of TRPM2 channels. We provide analytical and experimental evidence to support that the activation mode of TRPM2 channels by ADPR and Ca^2+^ evolved through at least three identifiable stages from a simple activation mode to an intermediate one and then to a complicated and coordinated one. We have identified permissive mutations which contribute to changes in the channel activation mode. We propose that the environmental oxygen level may facilitate the TRPM2 channels to acquire a more precisely regulated, coordinated activation mechanism. Our study could serve as a novel case for studying the evolution of other ion channels, receptors or proteins, and facilitate the understanding of how species adapted with their living environments from the perspective view of protein evolution.

## Supporting information

Main figures

## Acknowledgements

This work was supported by grants from the National Natural Science Foundation of China (82030108 and 31872796 to WY, 32071102 to PLY, 32000707 to CXG), the Zhejiang Provincial Natural Science Foundation of China (LD24H090004 and R16H090001 to WY, LQ20H160039 and LTY21H160003 to XZY, and LY19B020013 to PLY), National Major Scientific and Technological Special Project for “Significant New Drugs Development” (2018ZX09711001-004-005 to WY), Zhejiang Association for Science and Technology Talent Cultivation Project (CTZB-2020080127 to PLY), the East-West Cooperation Project (2019BFH02003 to WY), and the MOE Frontier Science Center for Brain Science & Brain-Machine Integration, Zhejiang University.

## Author contributions

W.Y. and C.M. designed the project; C.M., C.C and P.L.Y collected the datasets; C.M., Y.P.L and C.Y.Z. conducted the experiments and data analysis; C.M. and W.Y. drafted the manuscript; L.-H.J., W.Y., C.M., G.J.Z., J.-H.L. and F.Y. revised the manuscript. N.H., X.C.L. and J.A.W. assisted experiments and data analysis during the revision of the manuscript.

## Supplementary Figures

**Figure S1.**
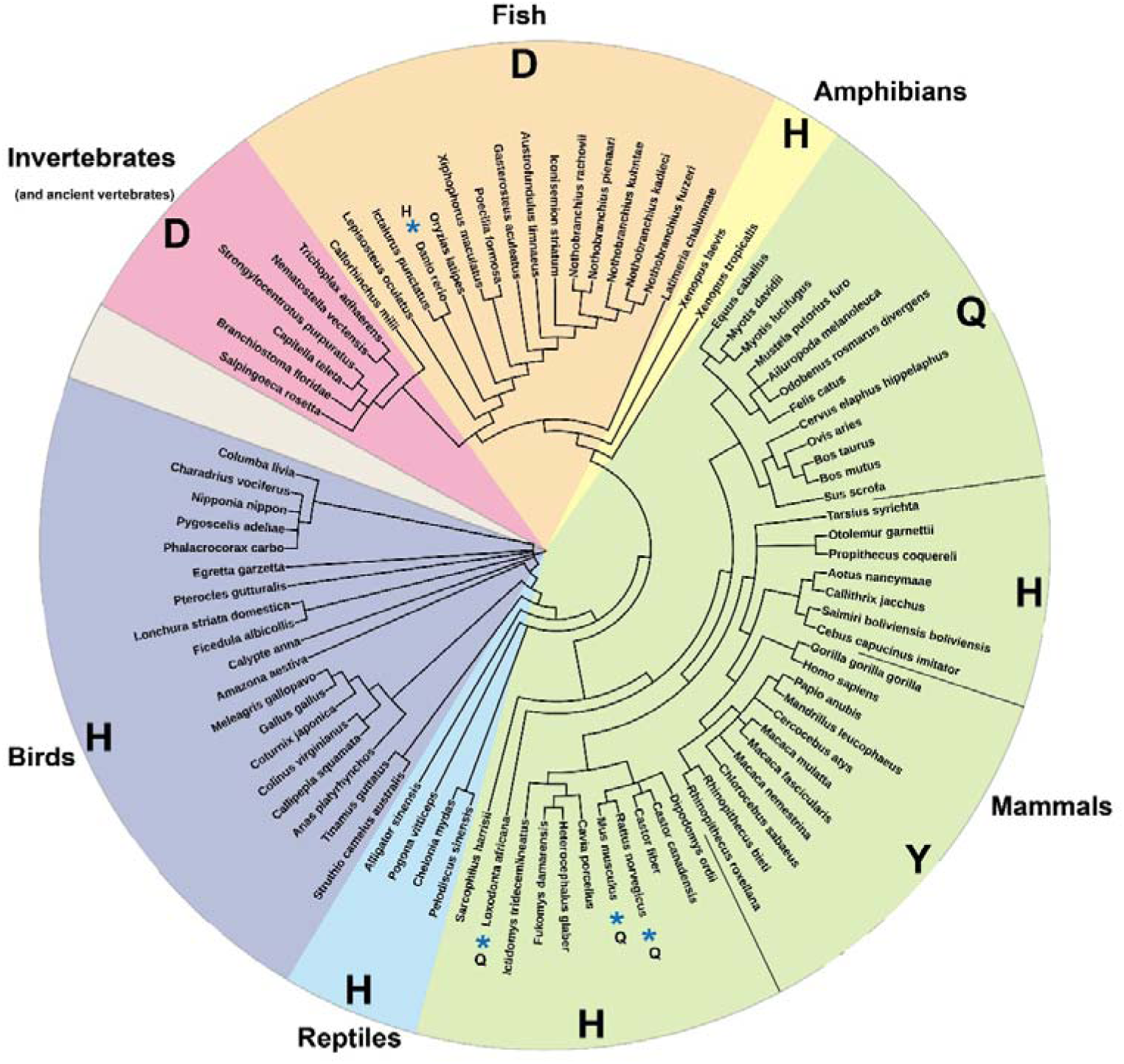
Maximum likelihood tree of TRPM2 channels. TRPM2 and TRPM2-like protein sequences of 89 species, from Unicellular flagellates, Flat animals, Jellyfish/anemones, Worms, Sea stars/urchins, Lancelets, Cartilaginous fishes, Bony fishes, Amphibians, Reptiles, Birds and Mammals. The amino acids corresponding to Y1349 in hsTRPM2 were labeled for each taxa, with blue * indicating the exceptional (i.e. “D” for *Danio rerio*, “Q” for *Loxodonta africana*, *Mus musculus* and *Rattus norvegicus*.)

**Figure S2.**
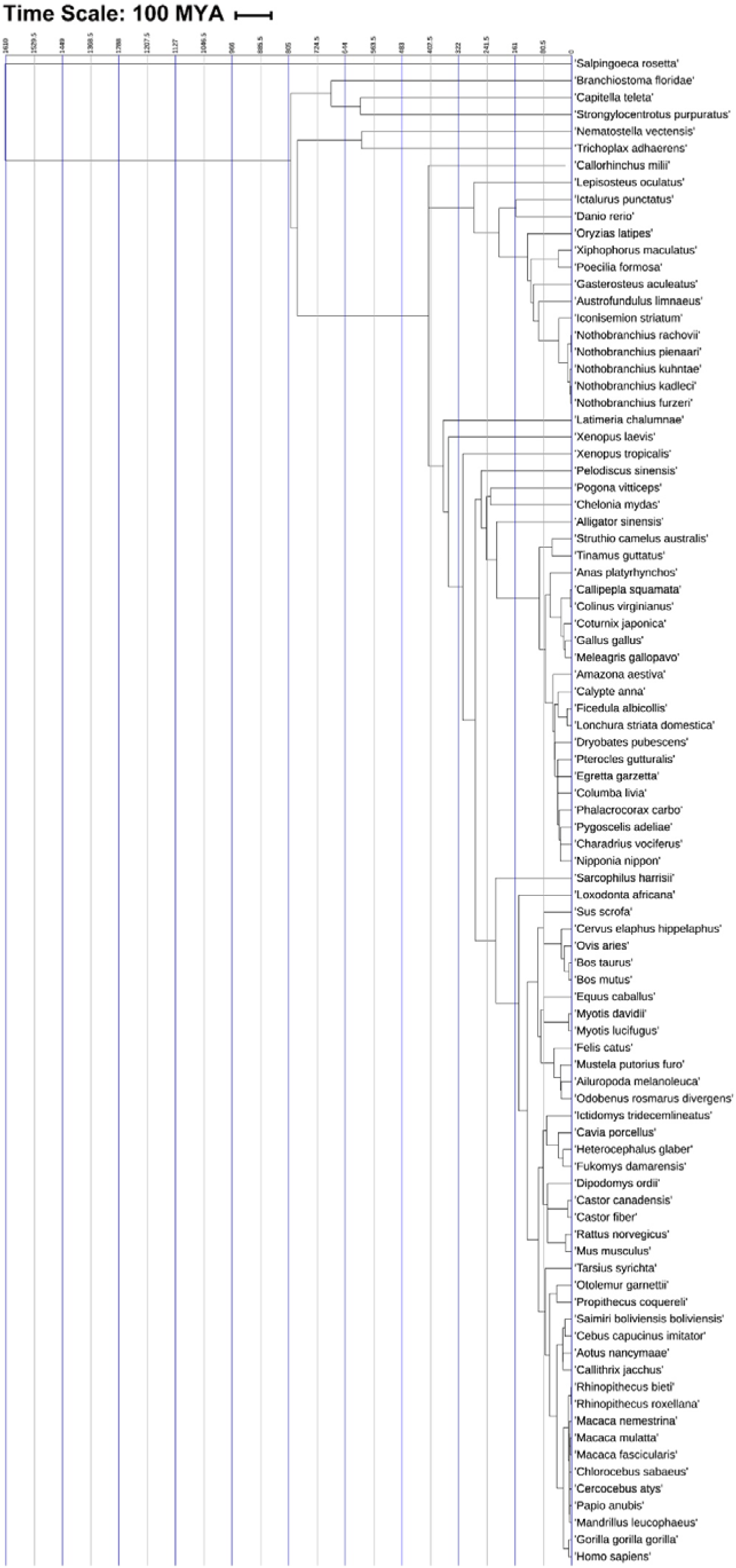
The original tree indicating the divergence times of TRPM2 channels. TRPM2 and TRPM2-like protein sequences were collected for 89 species exactly the same as indicated in Figure S1. The molecular “clock” was calibrated with constraints as listed in section 2.3, which were (1) Ancient: *Branchiostoma floridae* and *Strongylocentrotus purpuratus*, median time 627 MYA, (2) Cartilaginous fishes and Amphibian: *Callorhinchus milii* and *Xenopus laevis*, median time 465 MYA, (3) Amphibian and Reptile: *Xenopus laevis* and *Chelonia mydas*, median time 351.7 MYA, (4) Among Fish: *Oryzias latipes* and *Danio rerio*, median time 224 MYA, (5) Birds and Reptile: *Columba livia* vs *Chelonia mydas*, median time 262 MYA, (6) Among Primates: *Cebus capucinus imitator* and *Homo sapiens*, median time 42.9 MYA.

**Figure S3.**
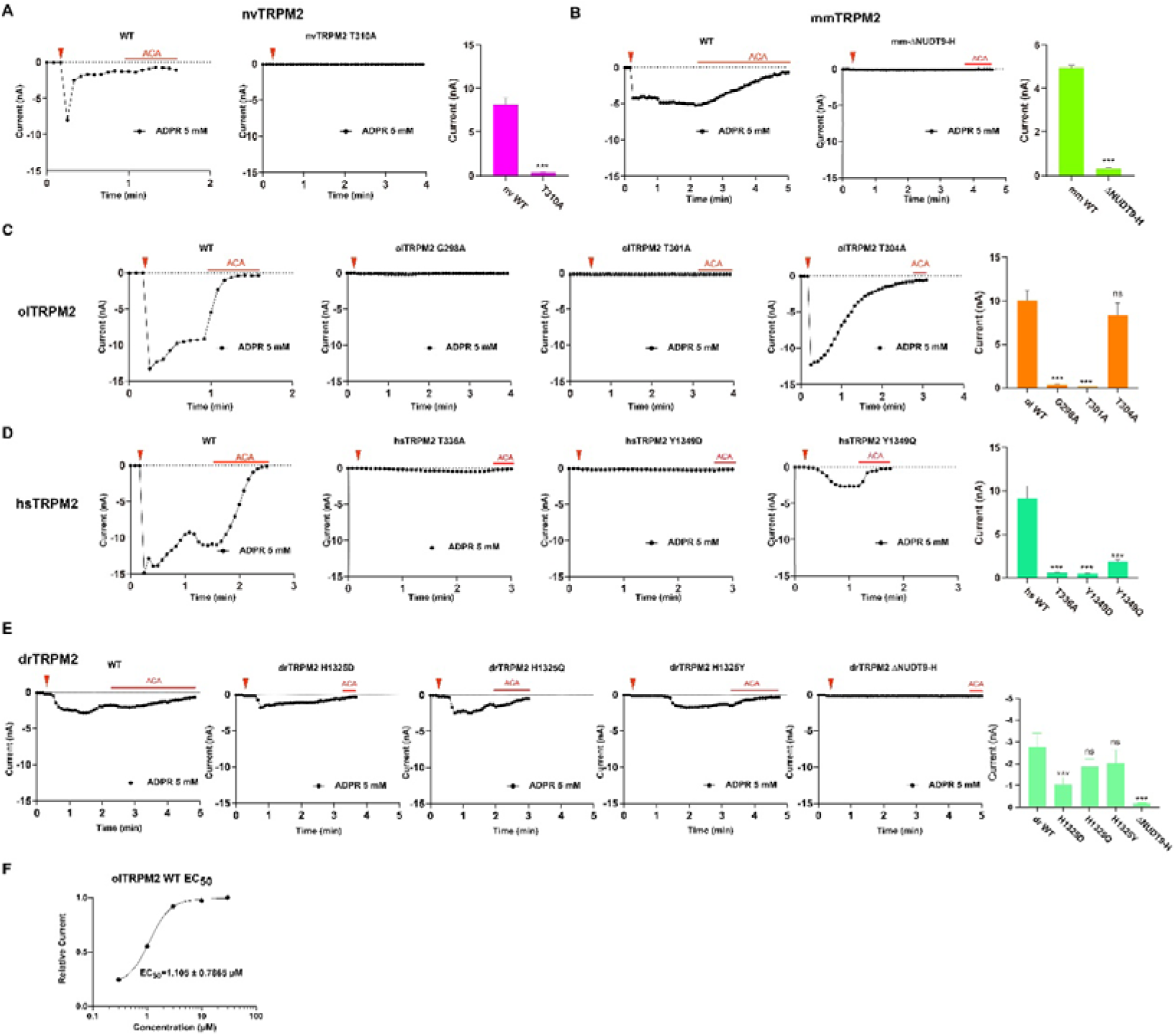
The activation of TRPM2 mutants by extreme ADPR concentration (5 mM). (A) to (E) Representative whole-cell patch recordings induced by 5 mM ADPR from HEK293T cells expressing WT or indicated mutant of nvTRPM2, olTRPM2, mmTRPM2, hsTRPM2 and drTRPM2. The red triangle in each panel indicates the time point at which whole-cell configuration was established. The data obtained by voltage ramps every 5 s; the red bars represent application of 20 μM N-(p-amylcinnamoyl) anthranilic acid. Summary of current densities were shown on the far right, data were expressed as mean ± starndard error of the mean (s.e.m), the number of cells examined in each case over three (n= 3-5). (F) The EC_50_ curve for the wild type olTRPM2.

**Figure S4.**
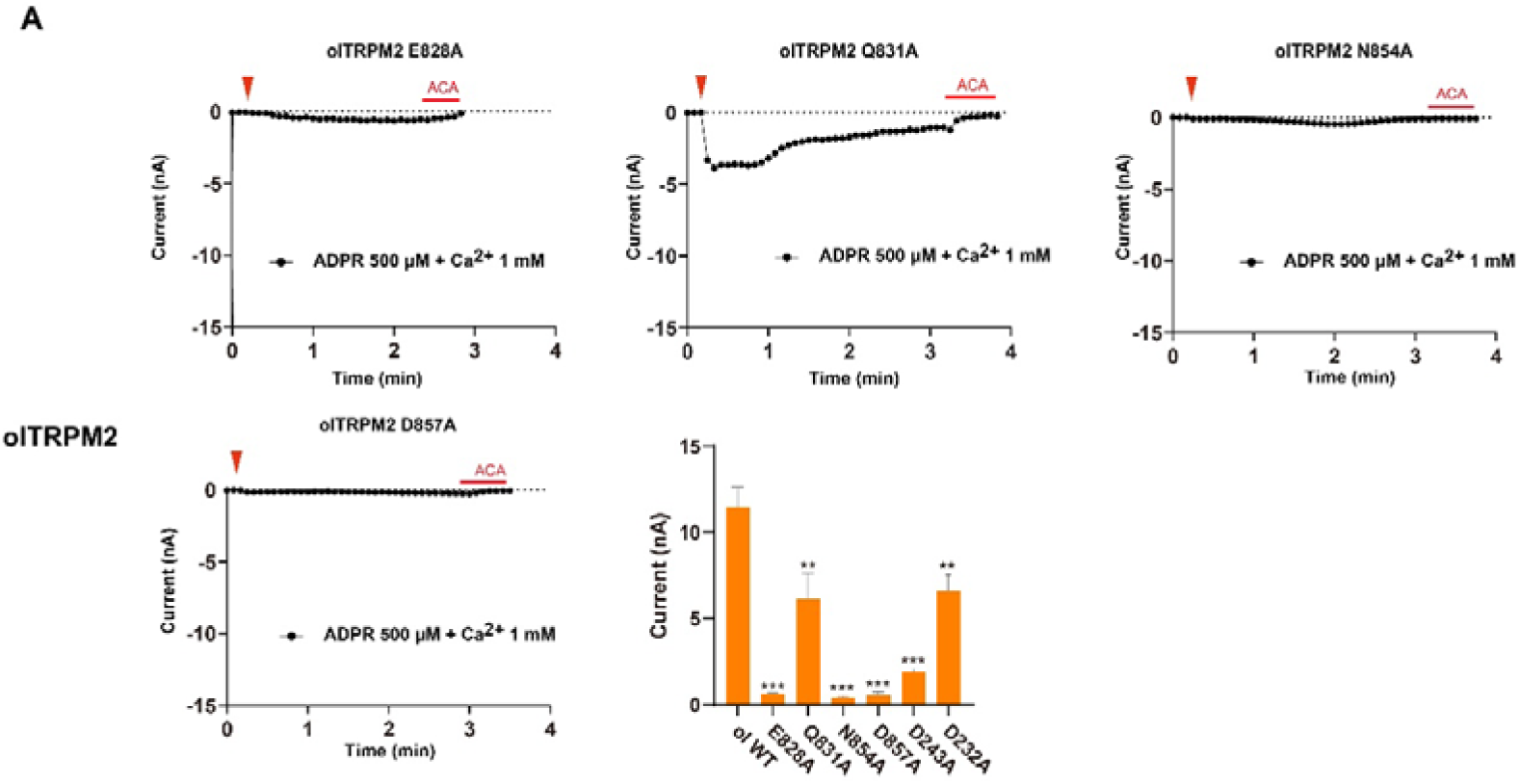
The activation of TRPM2 mutants by high Ca^2+^ concentration (1 mM). (A) Representative whole-cell patch recordings induced by 500 μM ADPR with additional 1 mM Ca^2+^ from HEK293T cells expressing WT or indicated mutant of olTRPM2. The red triangle in each panel indicates the time point at which whole-cell configuration was established. The data obtained by voltage ramps every 5 s; the red bars represent application of 20 μM N-(p-amylcinnamoyl) anthranilic acid. Summary of current densities were shown on the far right, data were expressed as mean ± starndard error of the mean (s.e.m), the number of cells examined in each case over three (n= 3-5).

**Figure S5.**
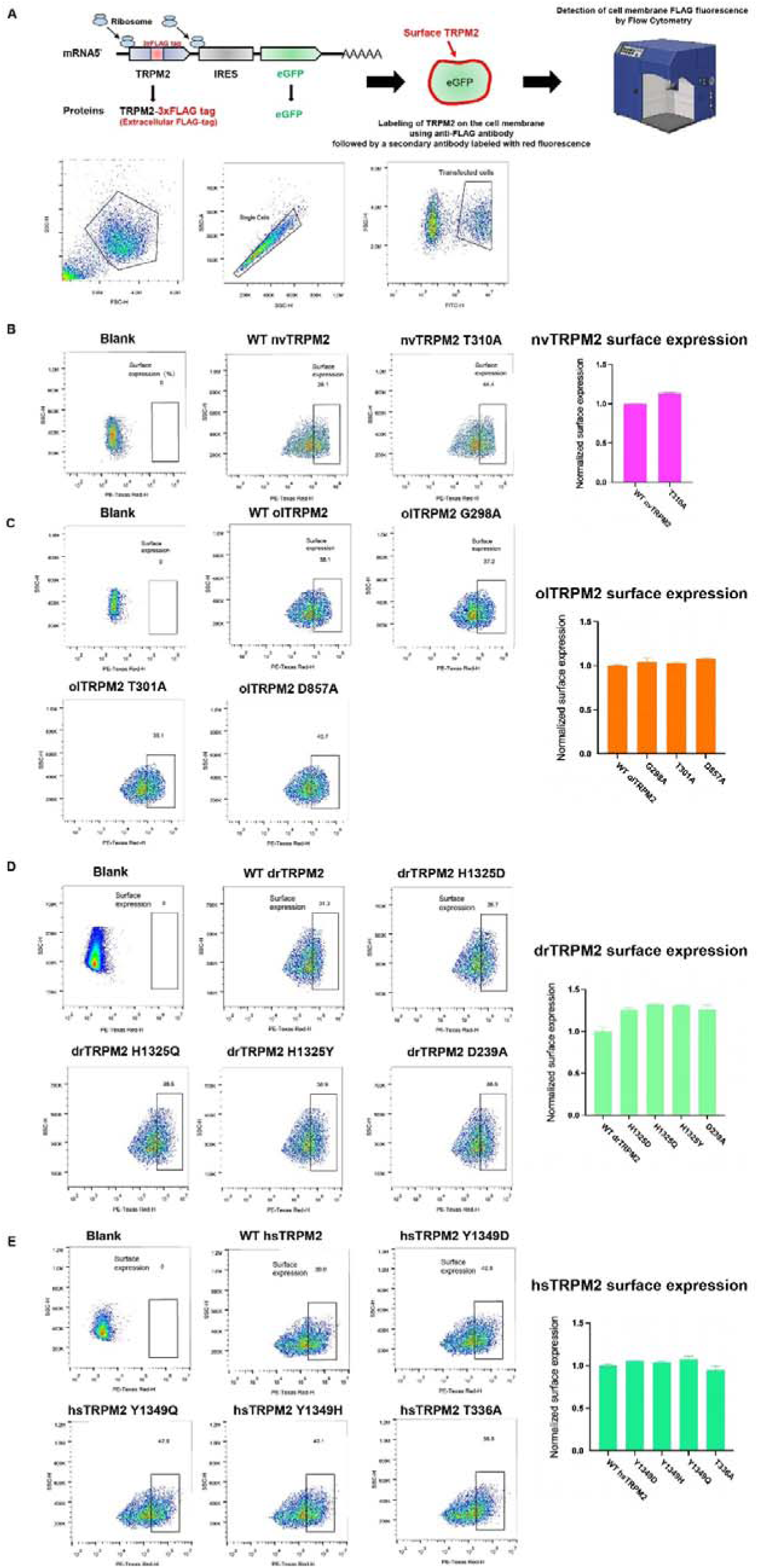
Membrane surface expression of WT and TRPM2 mutants. (A) Flow chart and a typical gating strategy for detecting surface expression of TRPM2 in HEK293T cells using flow cytometry. (B) to (E) Flow cytometry assays of membrane surface expression of WT TRPM2 and the indicated mutants. This includes WT and T310A of nvTRPM2 (related to Figure 3C and supplementary Figure 3A); WT and G298A, T301A and T304A of olTRPM2 (related to Figure 3C and supplementary Figure 3C); WT, Y1349D, Y1349Q, Y1349H and T336A of hsTRPM2 (related to Figure 3C and supplementary Figure 3D); and WT, H1325D, H1325Q, H1325H and D239A of drTRPM2 (related to Figure 6D)

**Figure S6.**
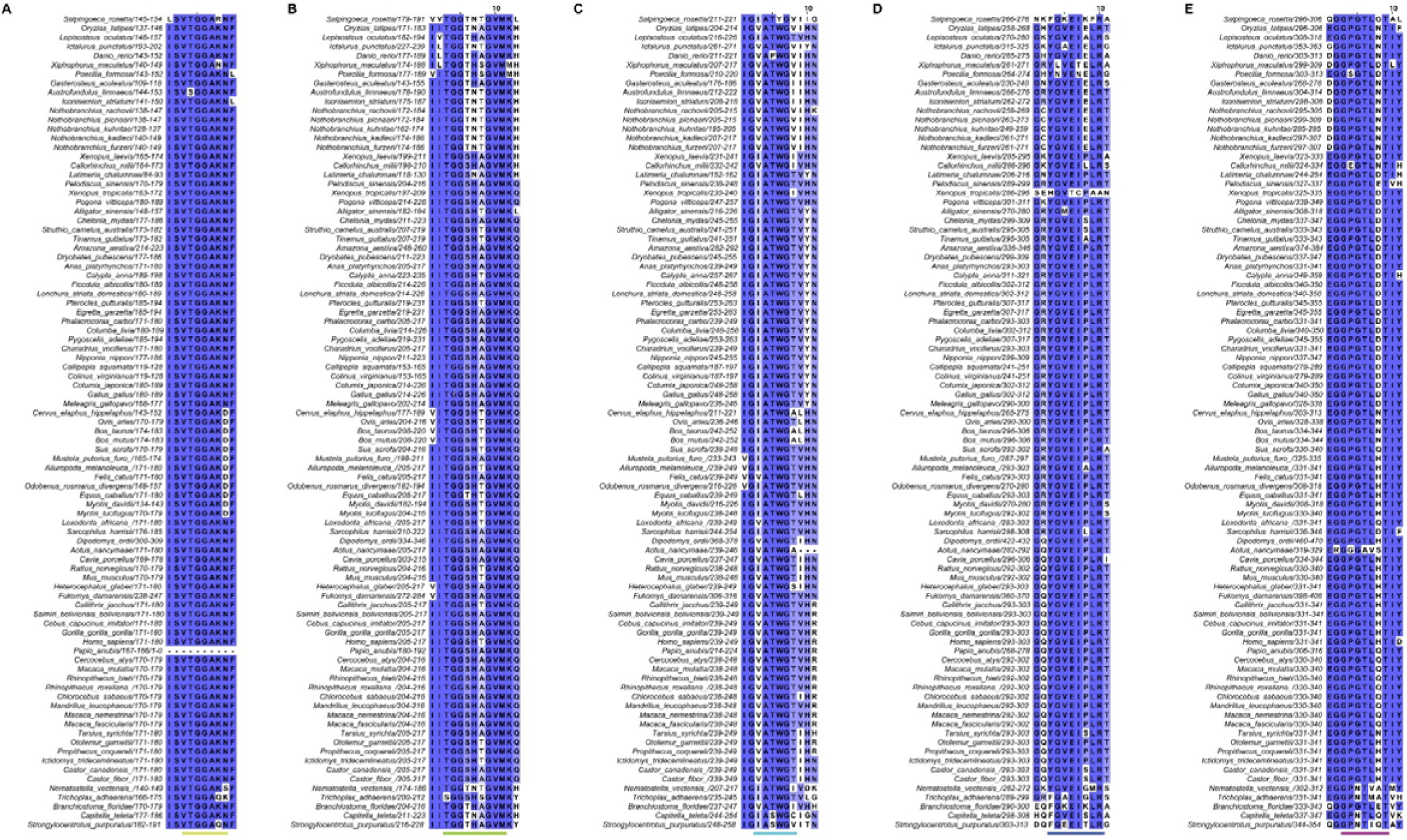
Sequence alignment of residues forming or around the N-termini ADPR binding pocket of TRPM2s from the 89-sequeces dataset.

**Figure S7.**
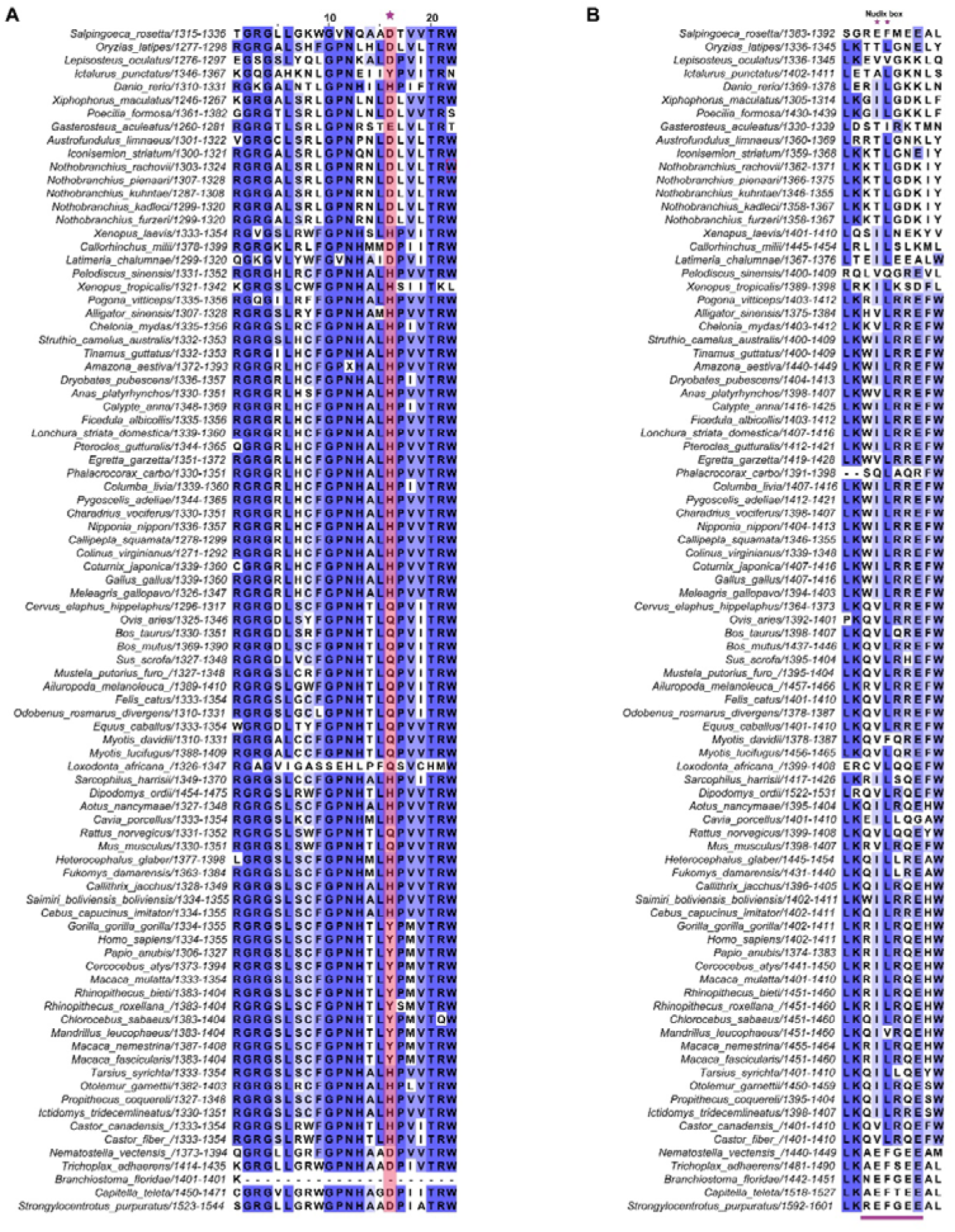
Sequence alignment of residues forming or around the C-termini ADPR binding pocket and the “Nudix box” of TRPM2s from the 89-sequeces dataset.

**Figure S8.**
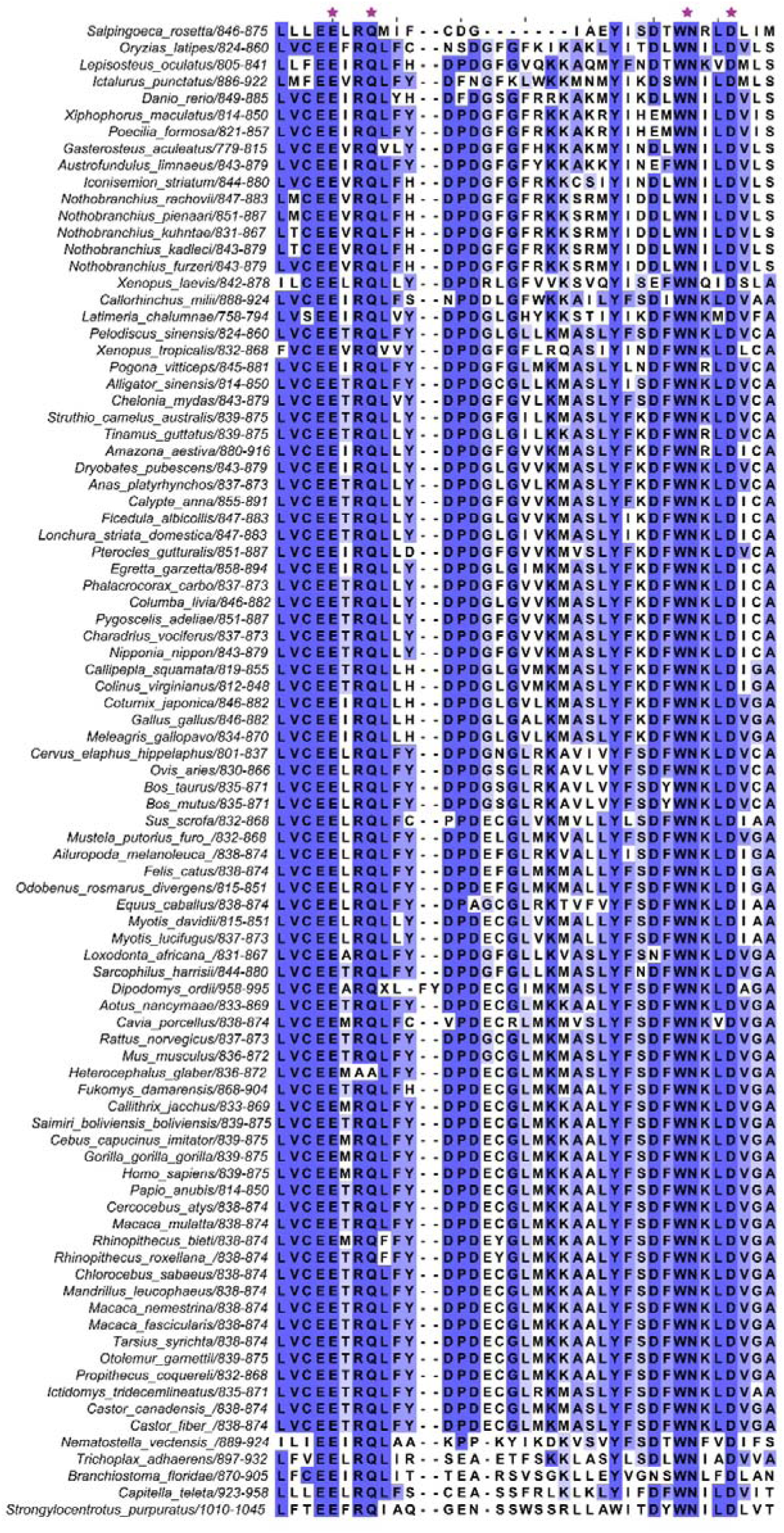
Sequence alignment of the S2-S3 residues involved in Ca^2+^ binding of TRPM2s from the 89-sequeces dataset.

**Figure S9.**
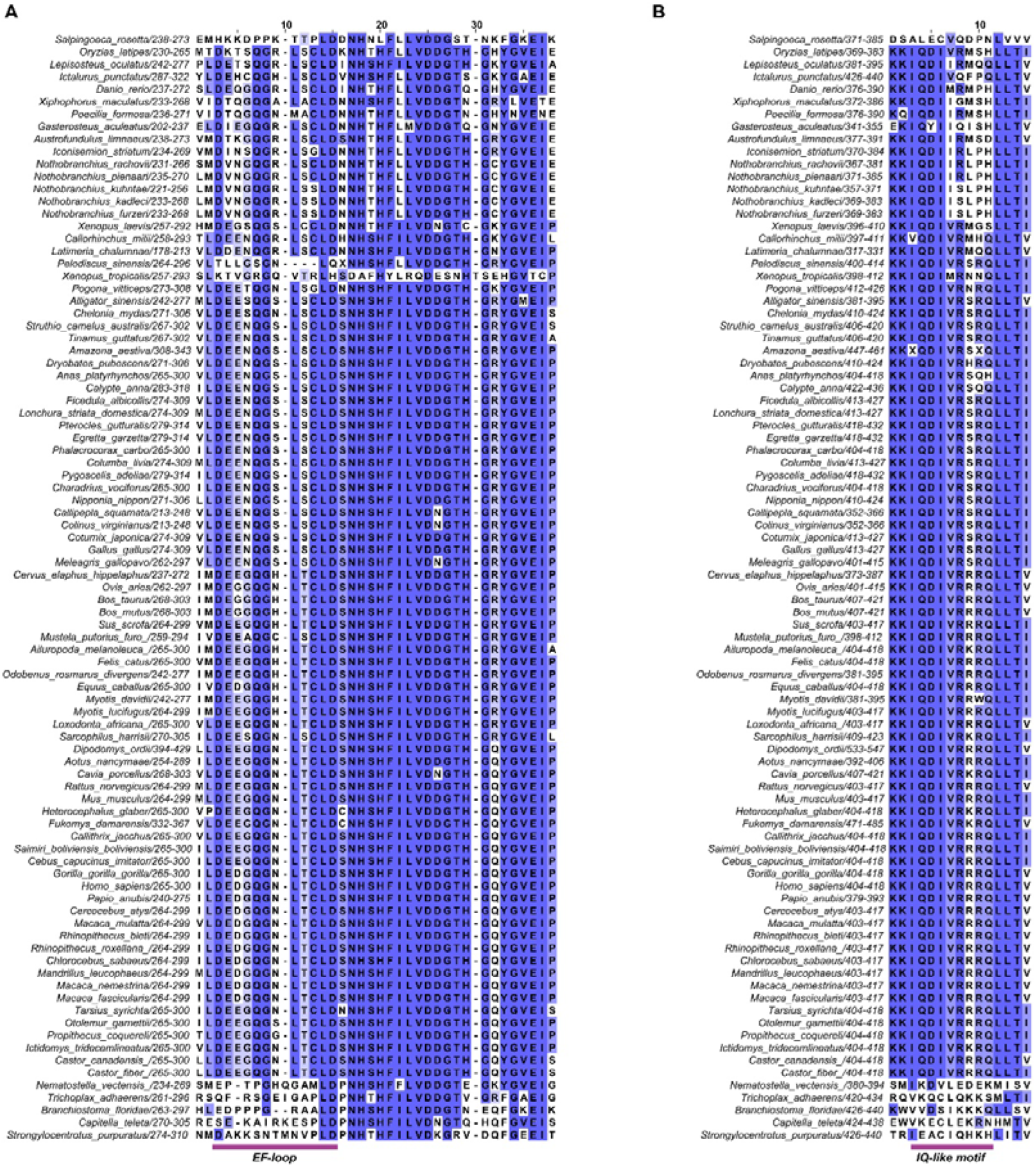
Sequence alignment of residues of or corresponds to EF-loop and IQ-like motif of TRPM2s from the 89-sequeces dataset.

