## Supplementary figures and images for "Evolutionary trajectory of TRPM2 channel activation by adenosine diphosphate ribose and calcium"

### Figure 1.jpg

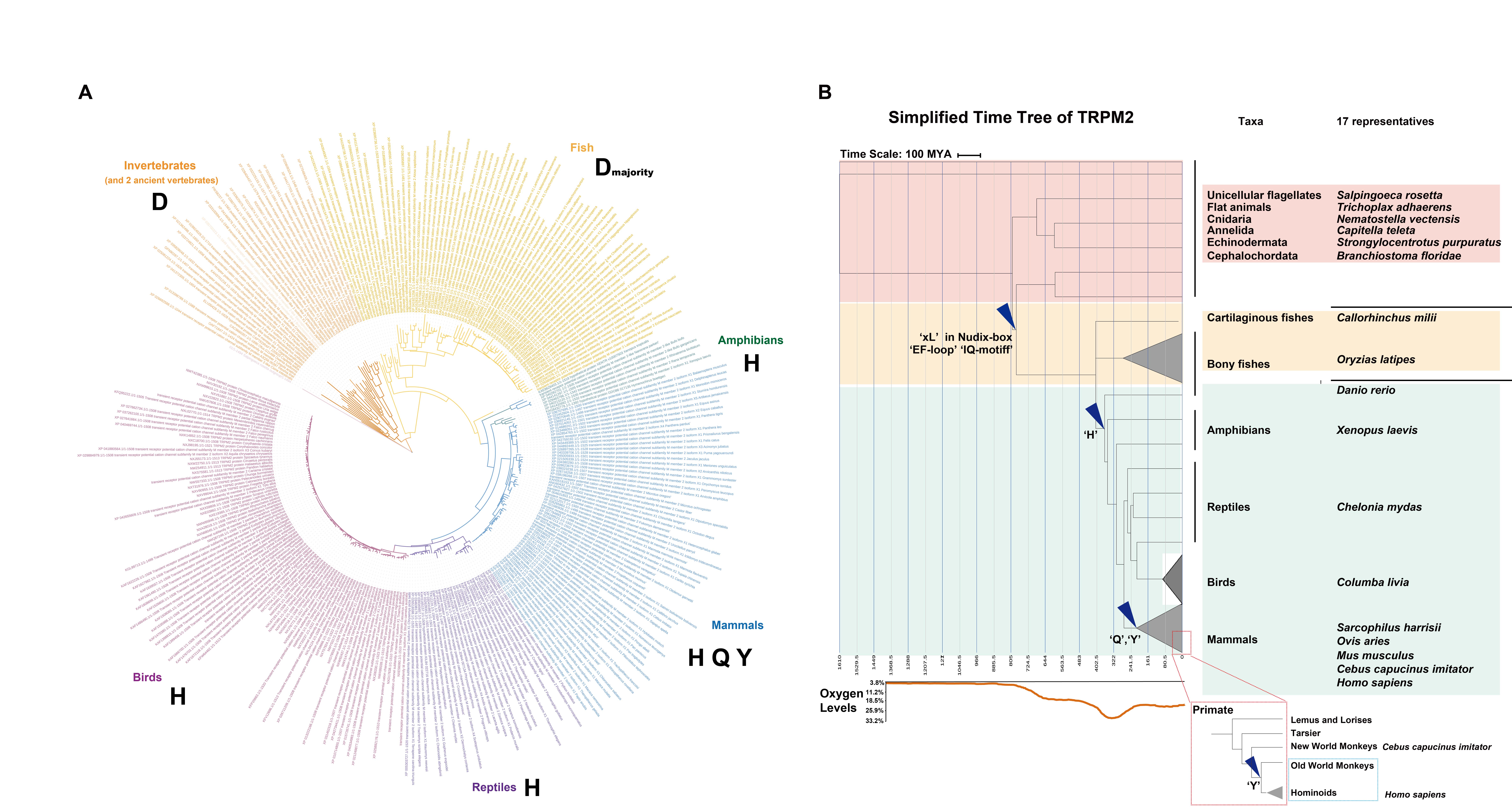

### Figure 2.jpg

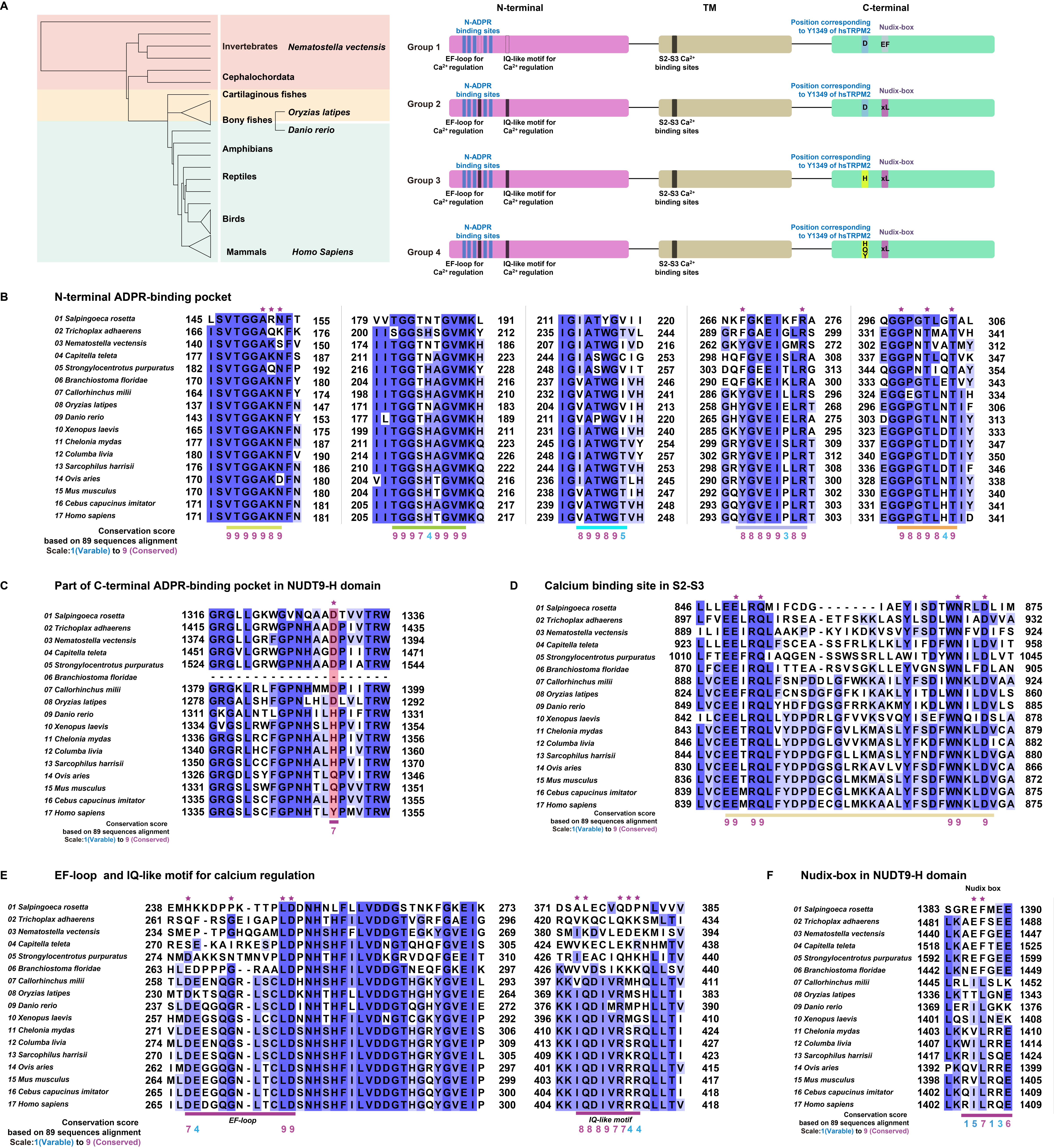

### Figure 3 N-ADPR.jpg

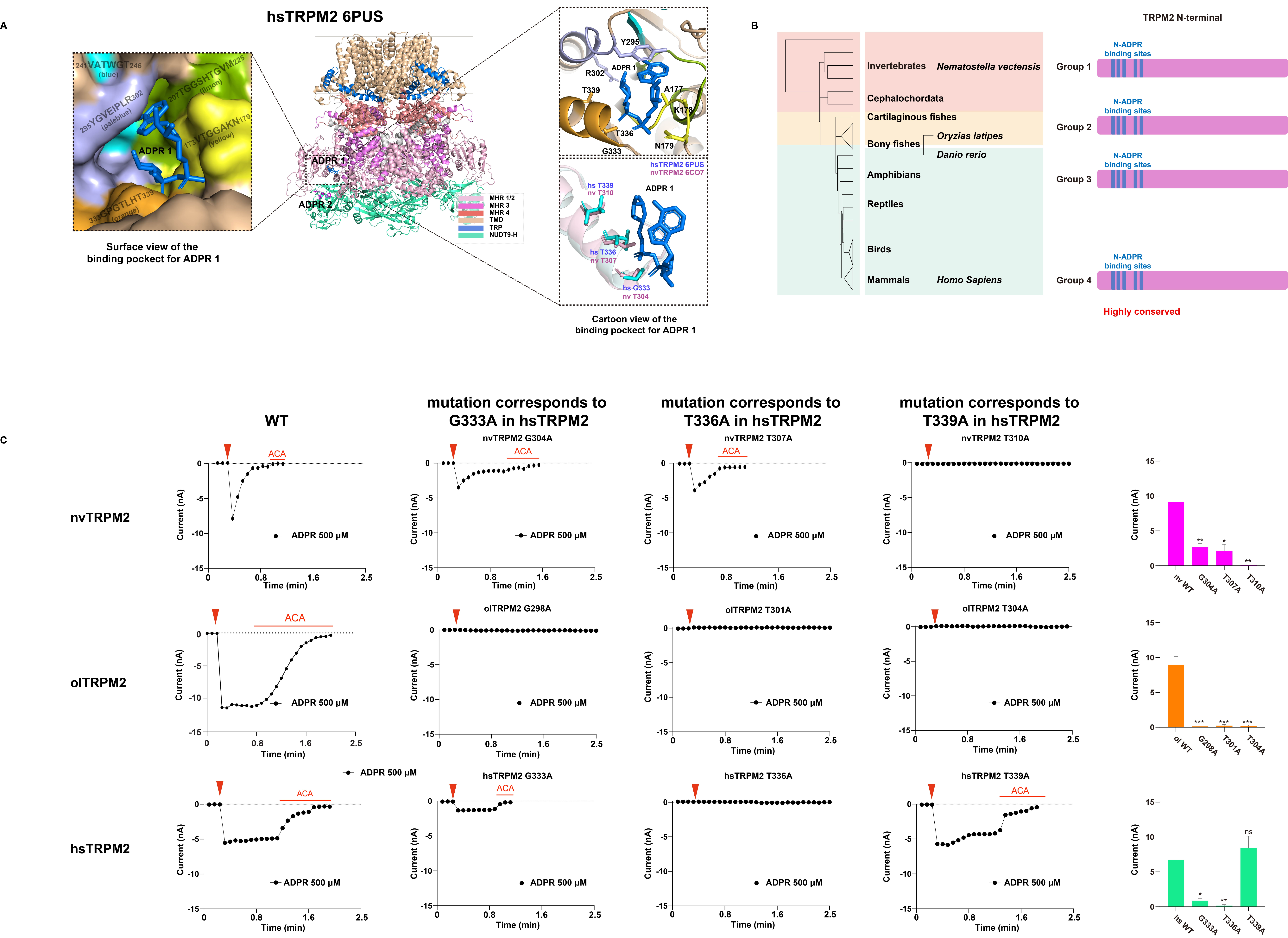

### Figure 4 Y1349.jpg

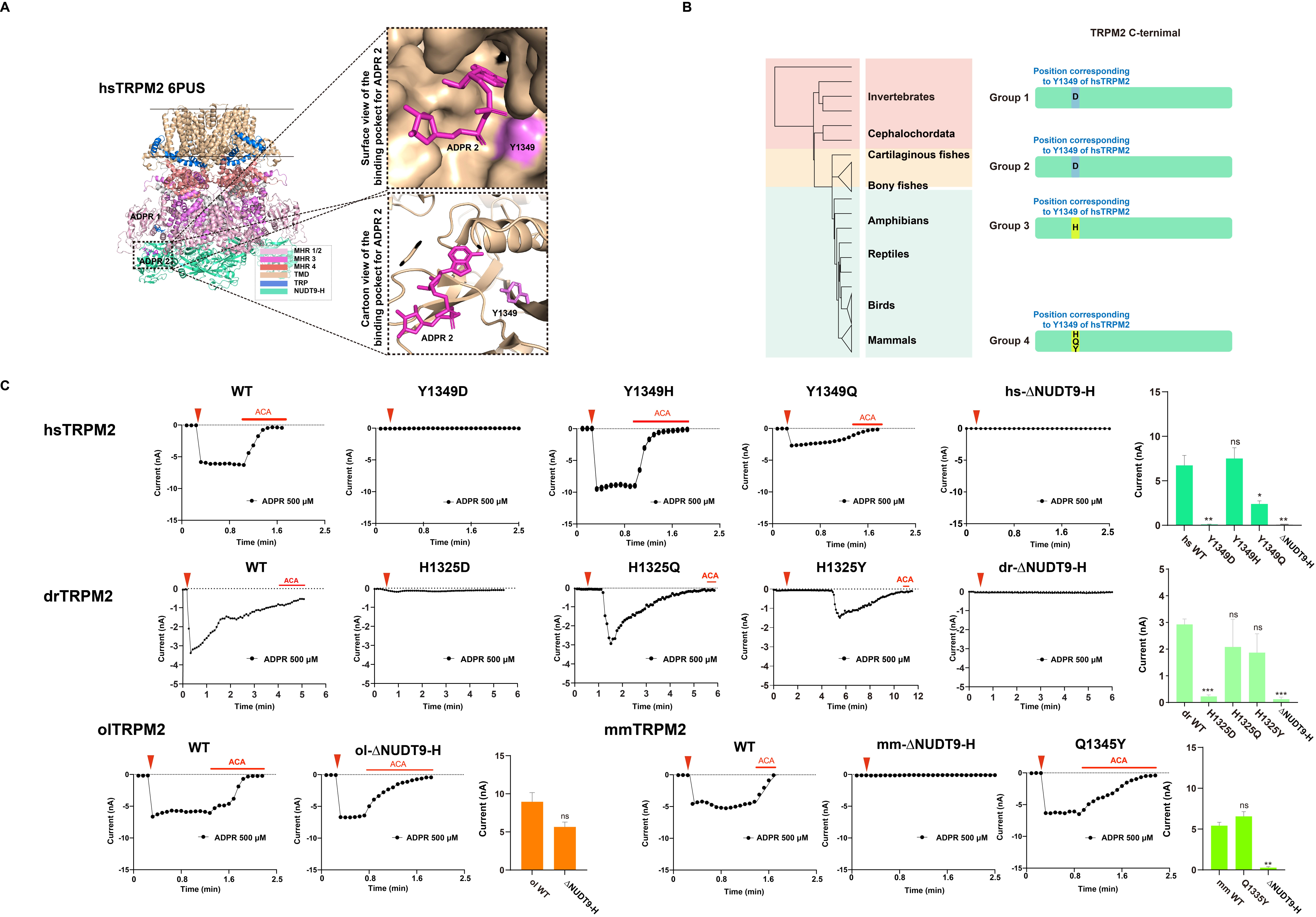

### Figure 5 s2-s3 binding site.jpg

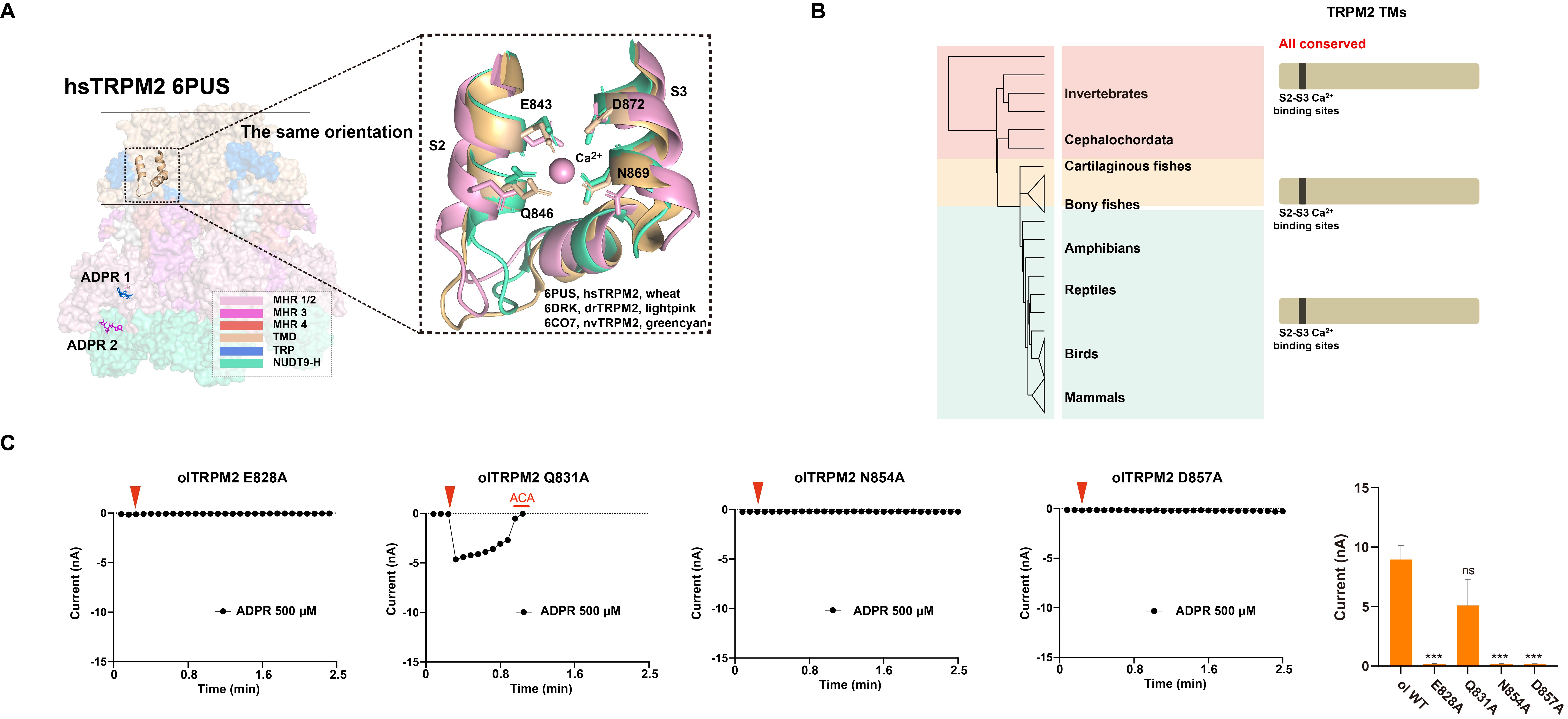

### Figure 6 EF1,EF2 and IQ.jpg

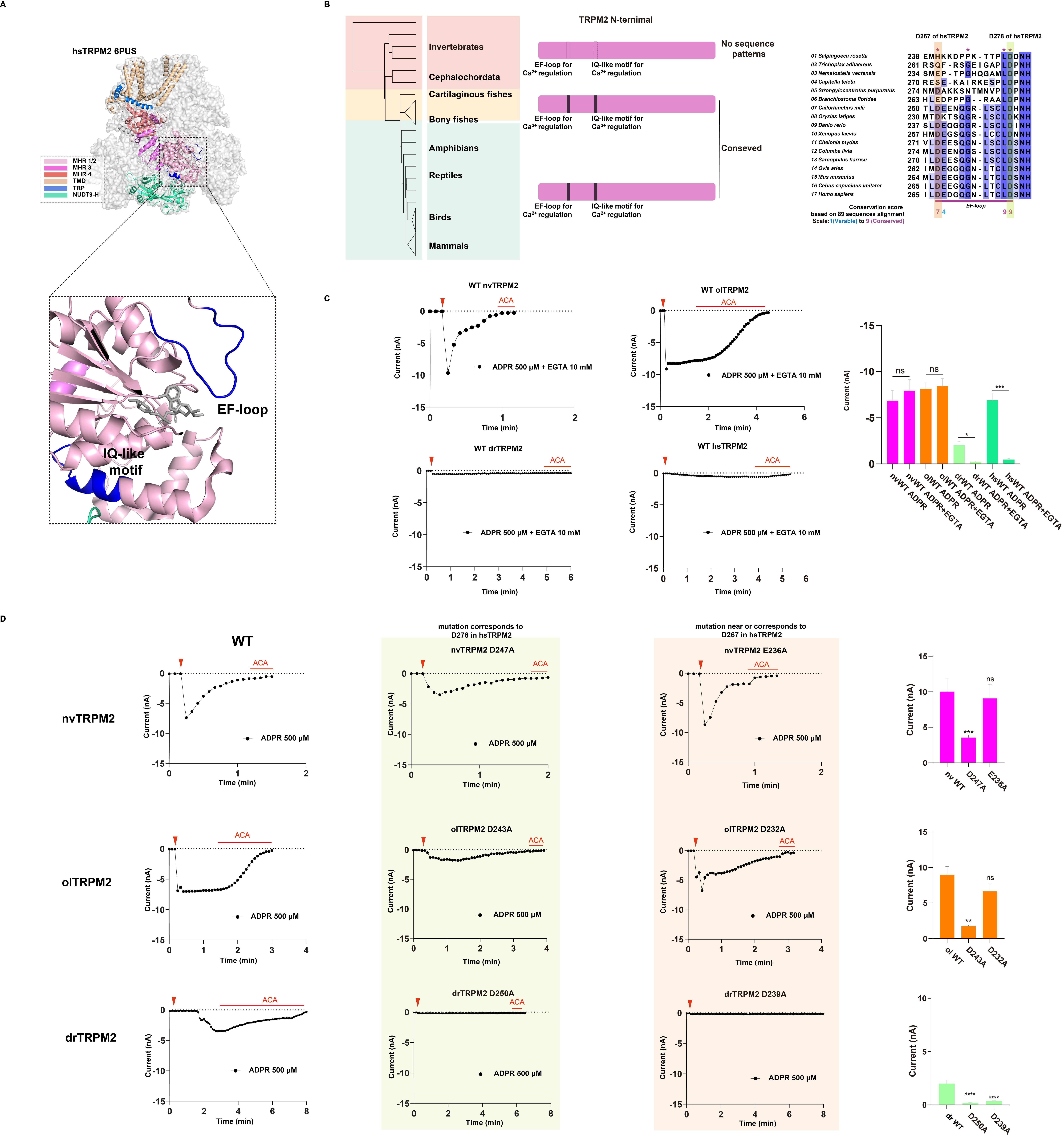

### Figure 7 models.jpg

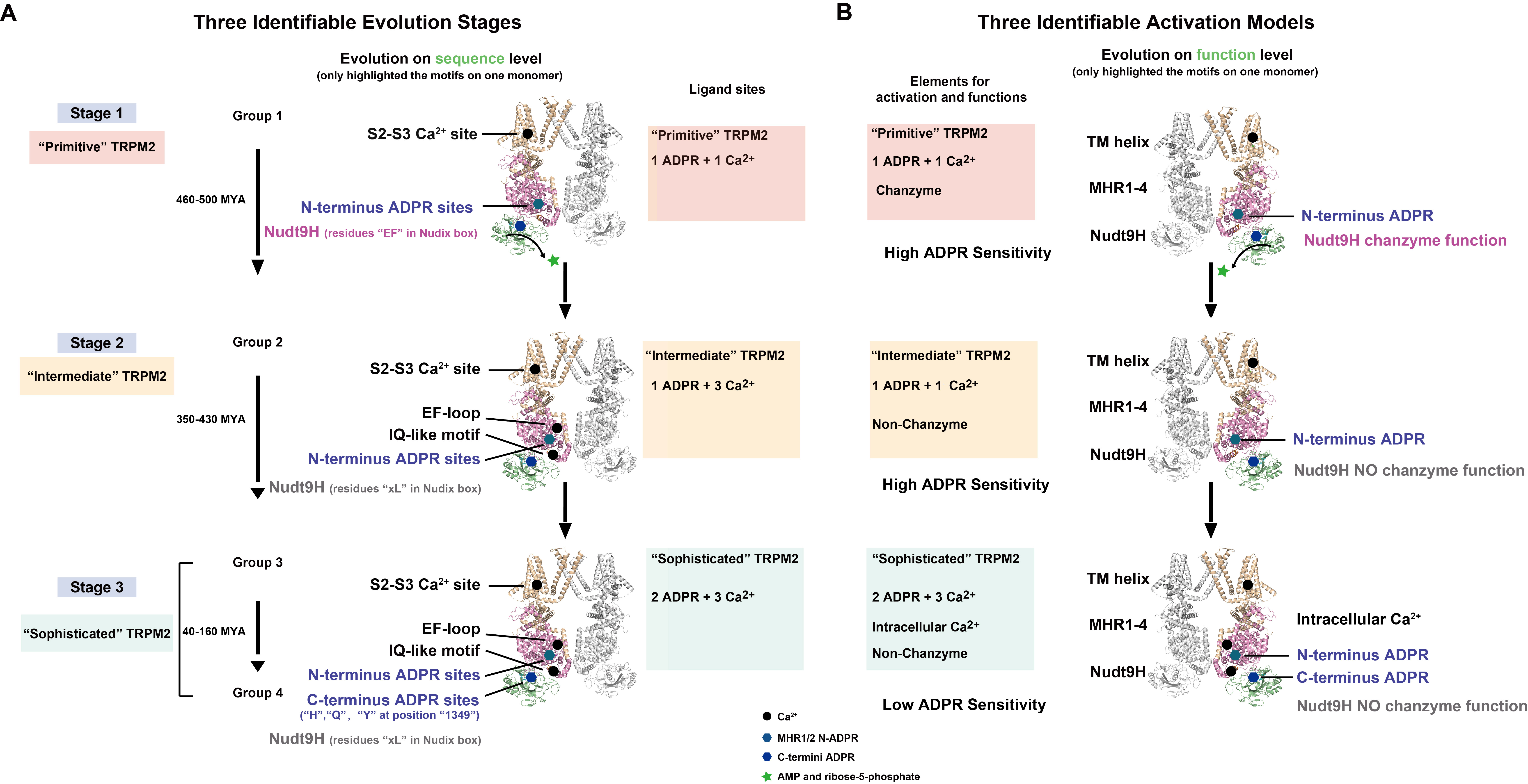

### supplementary Figure S1.jpg

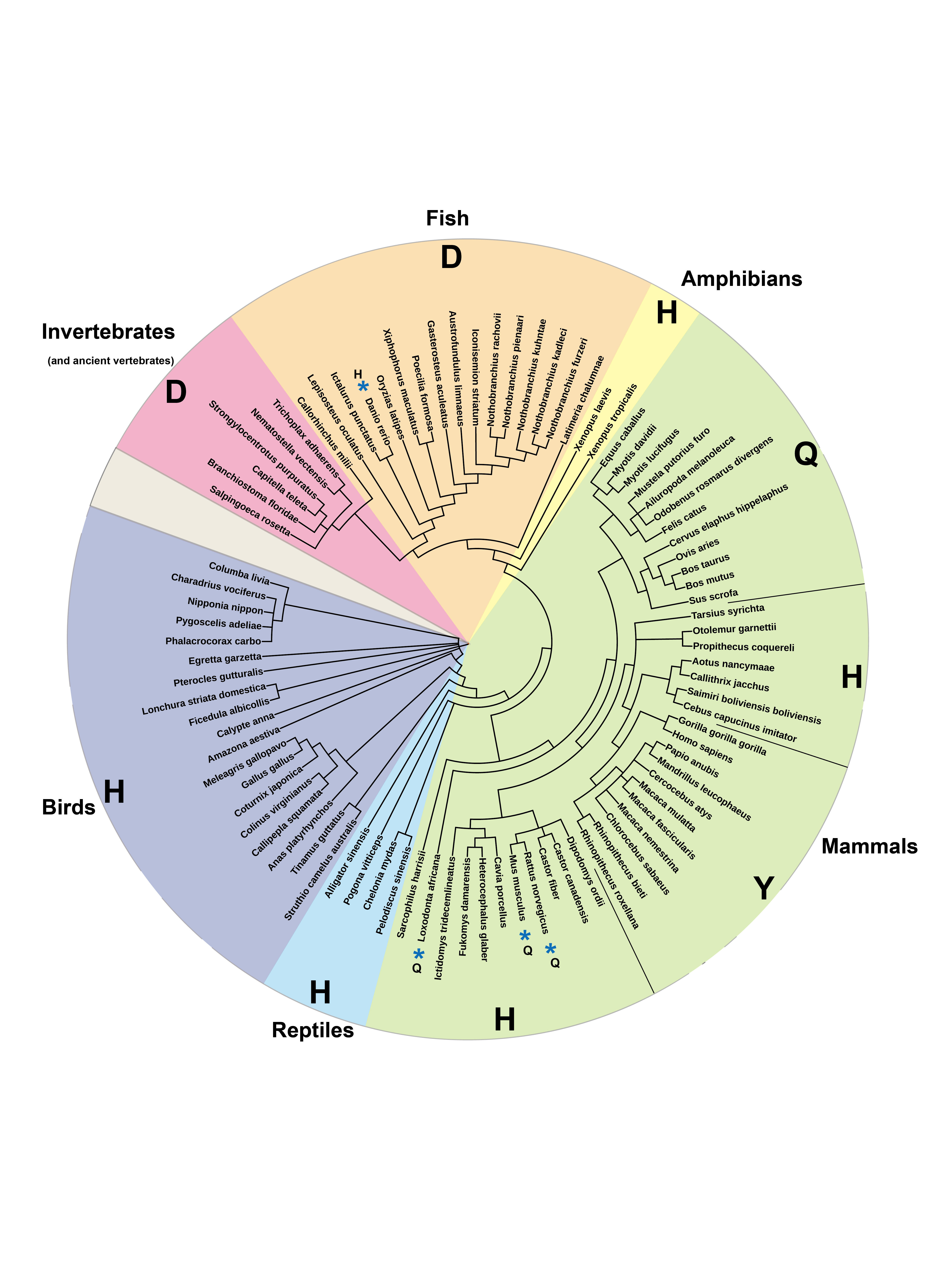

### supplementary Figure S2 Time tree.jpg

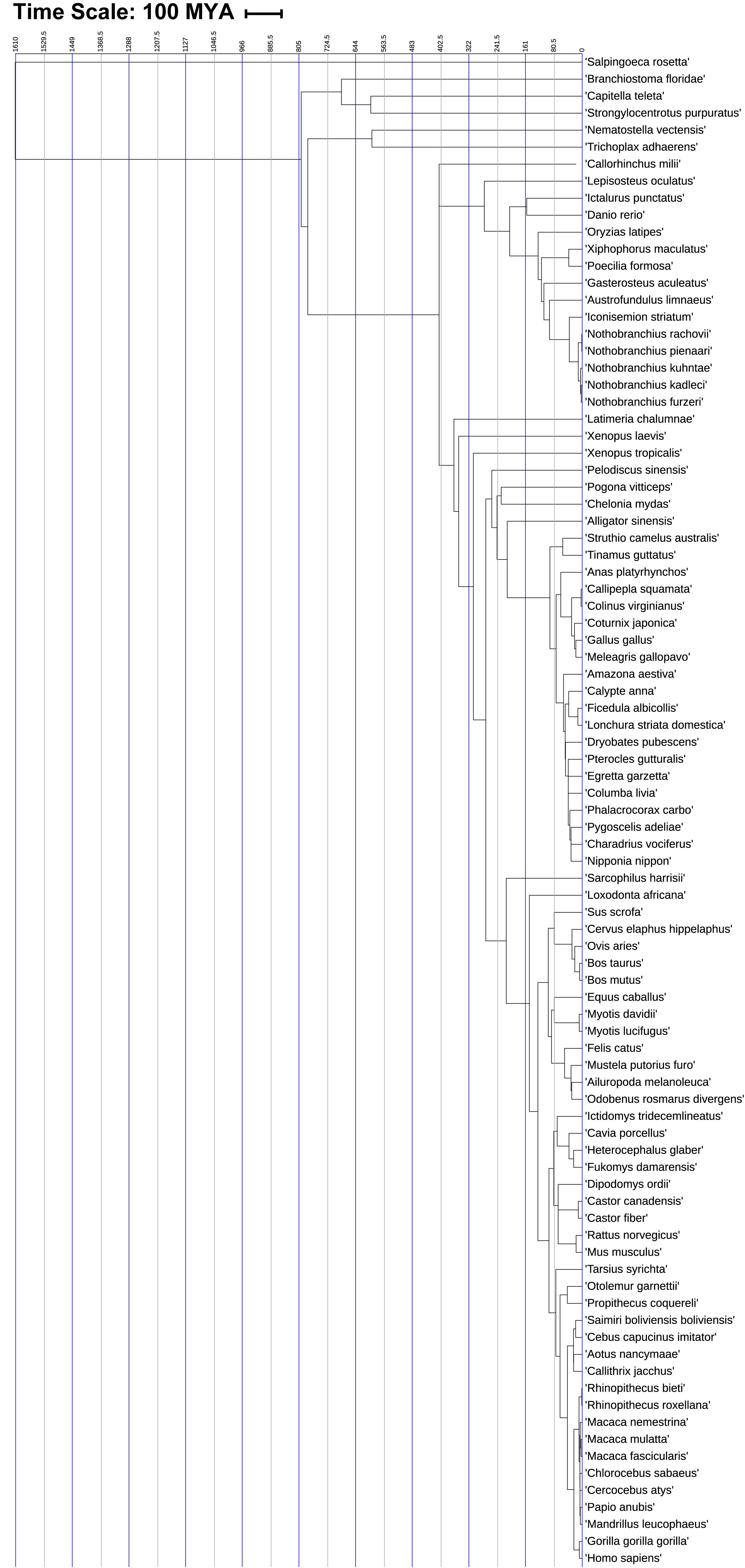

### supplementary Figure S3 ADPRhigh.jpg

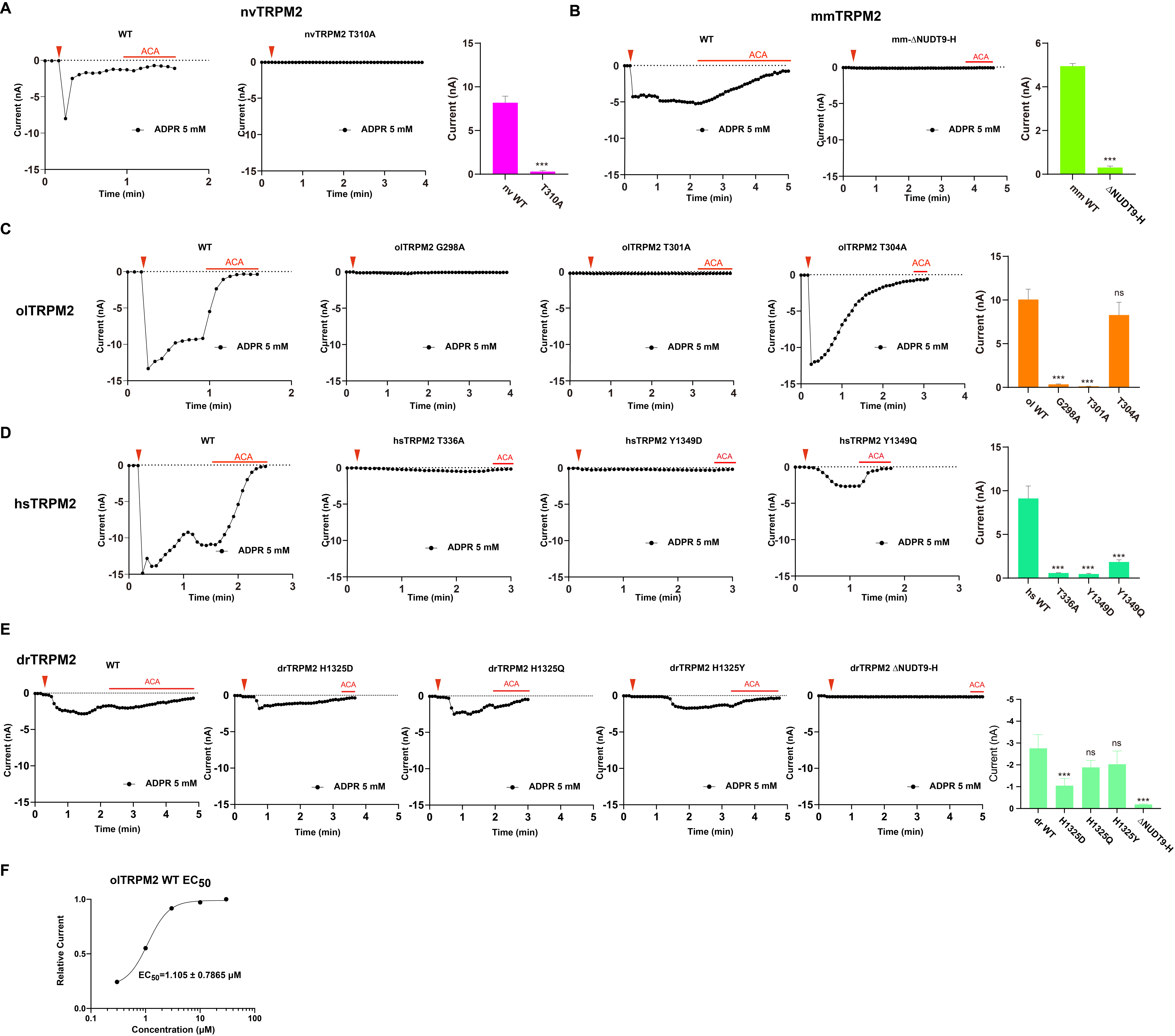

### supplementary Figure S4 Cahigh.jpg

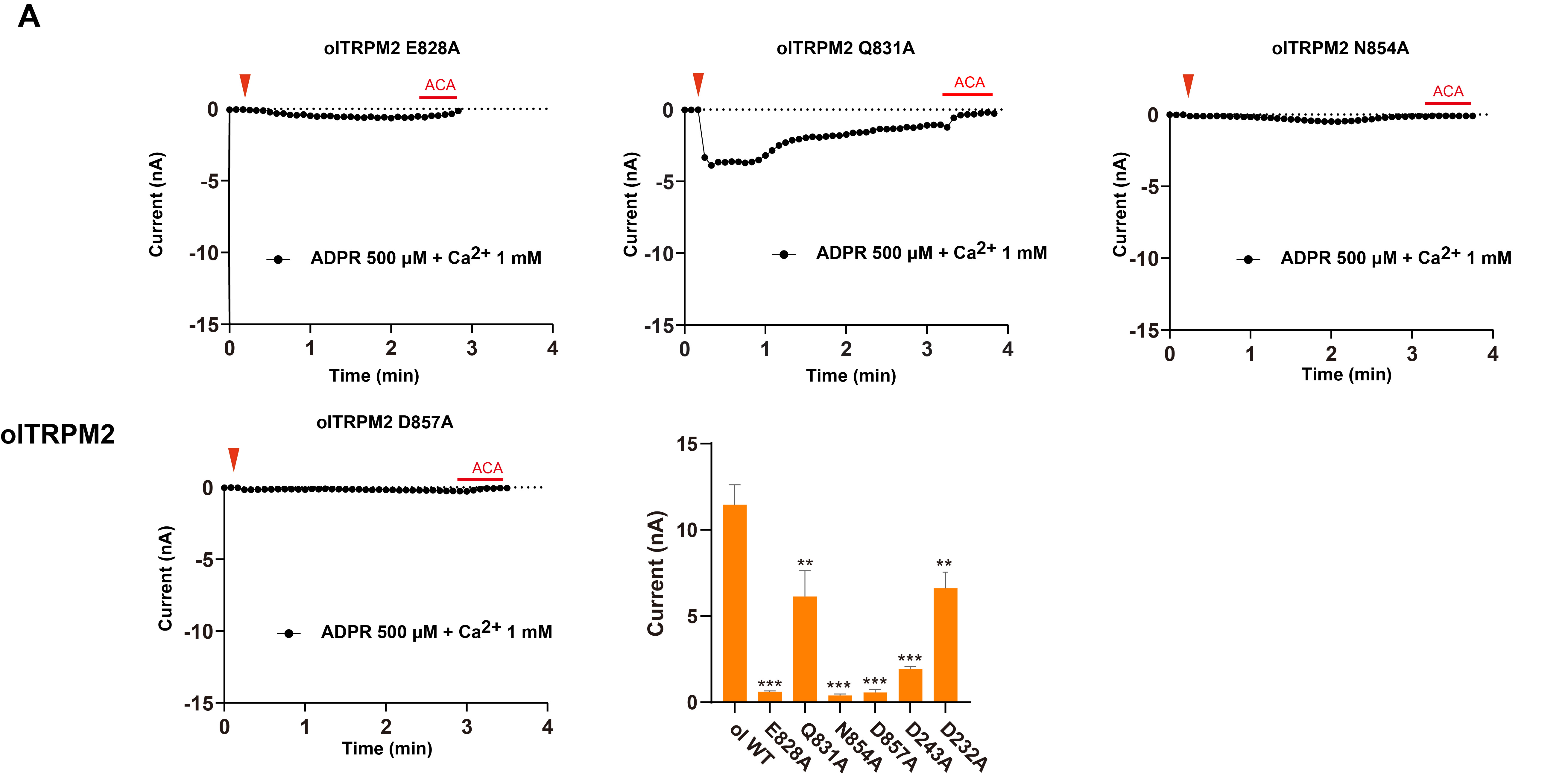

### supplementary Figure S5 surface expression.jpg

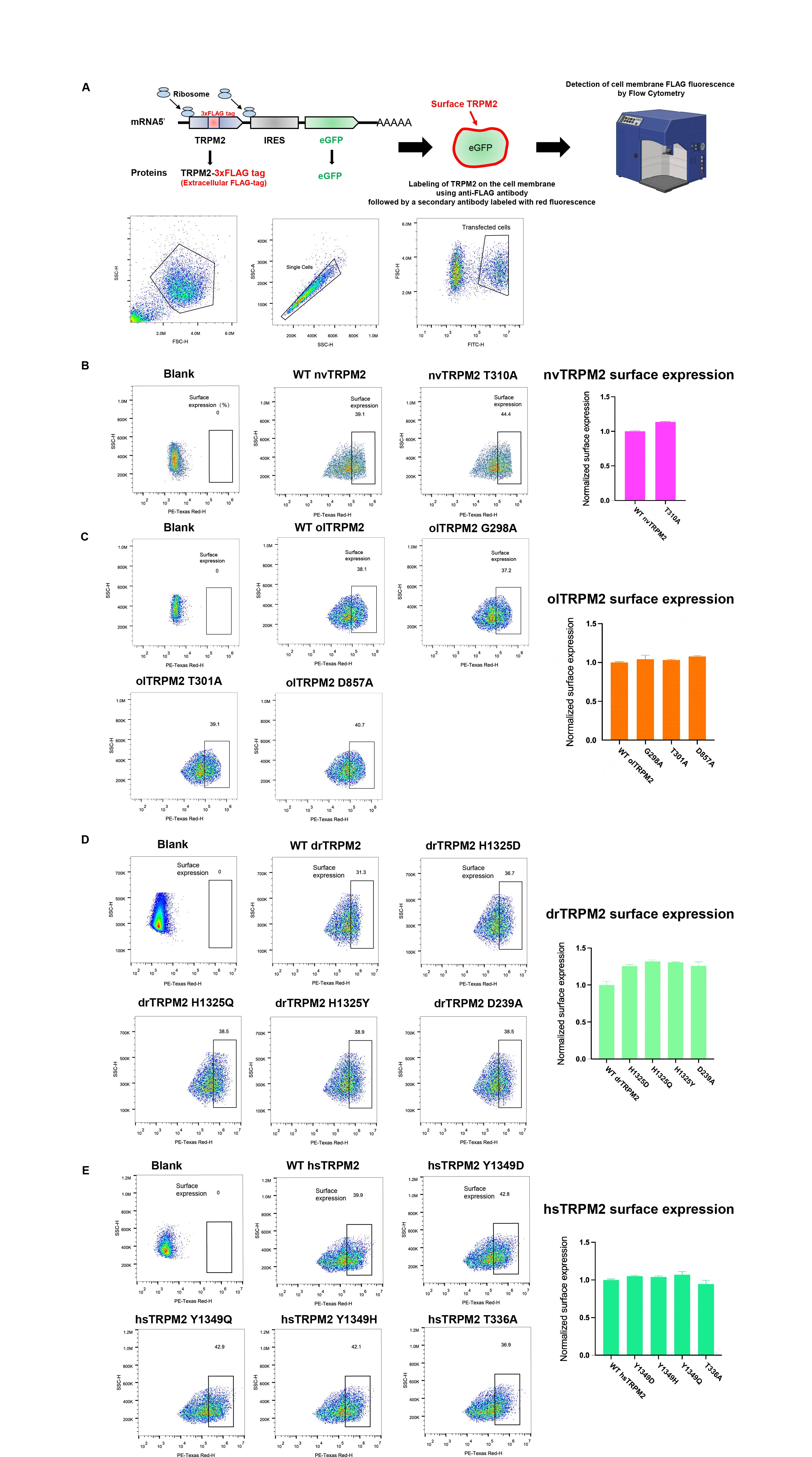

### supplementary Figure S6 89 N-pocket.jpg

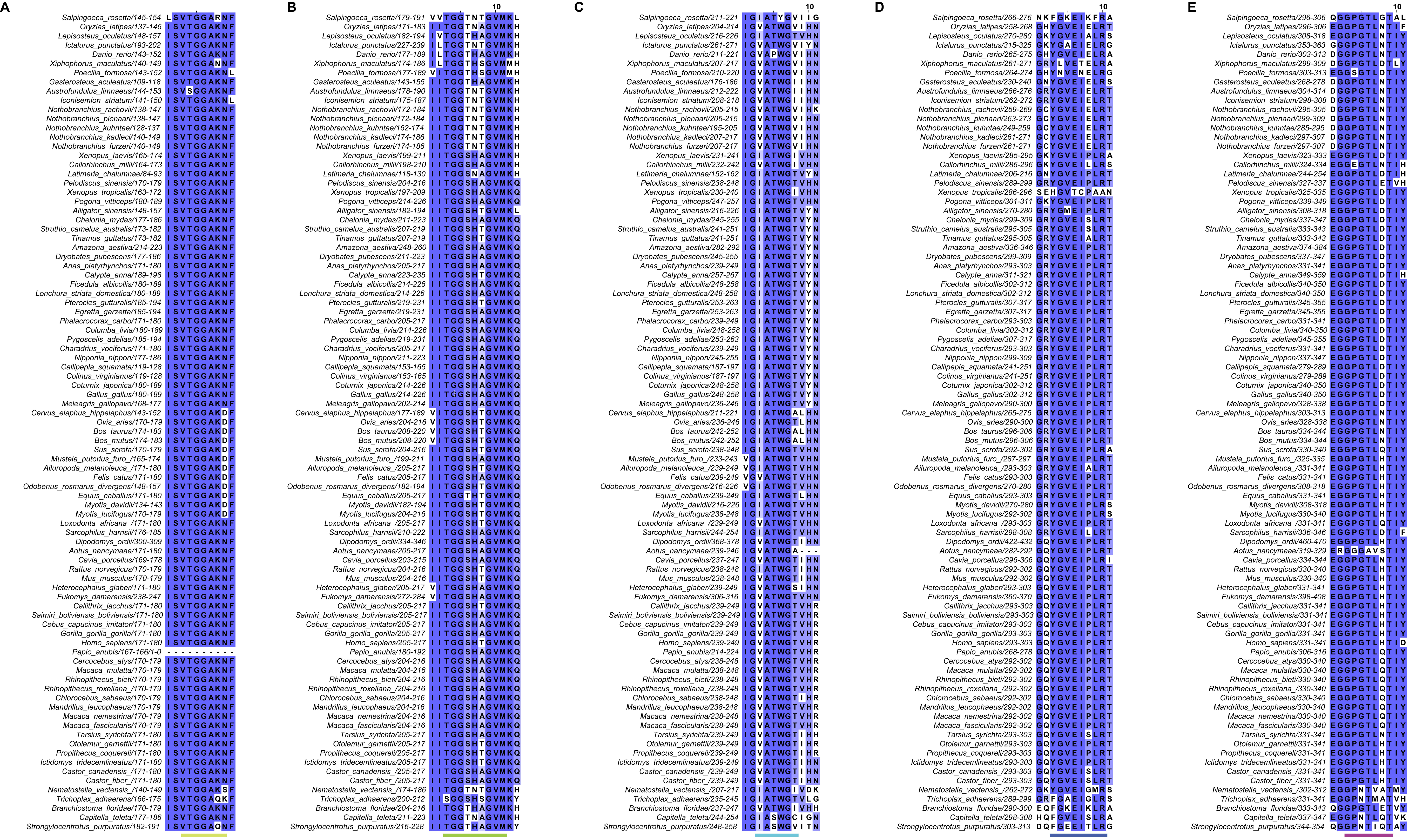

### supplementary Figure S7 89 Y1349-01.jpg

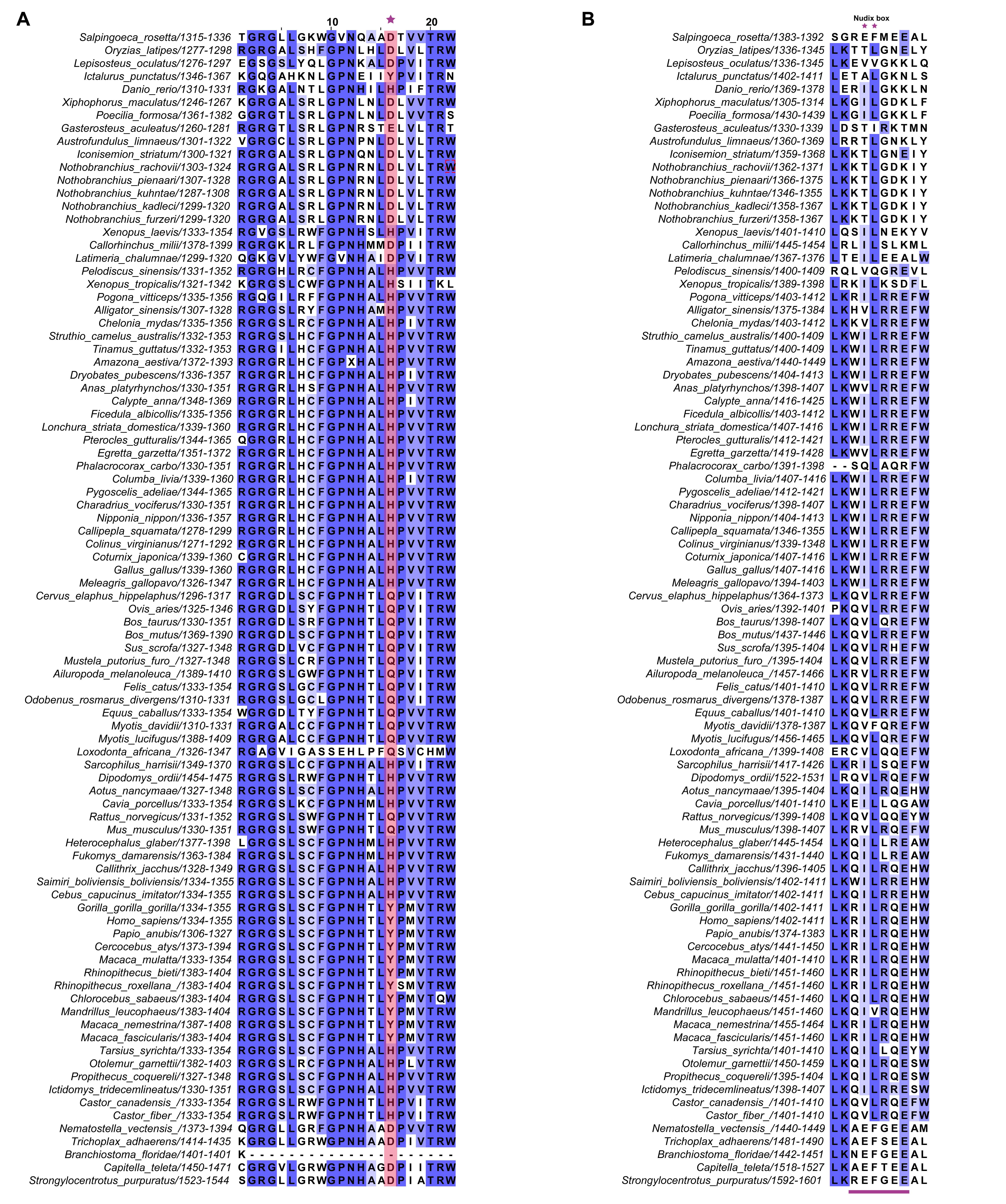

### supplementary Figure S8 89 S2-S3.jpg

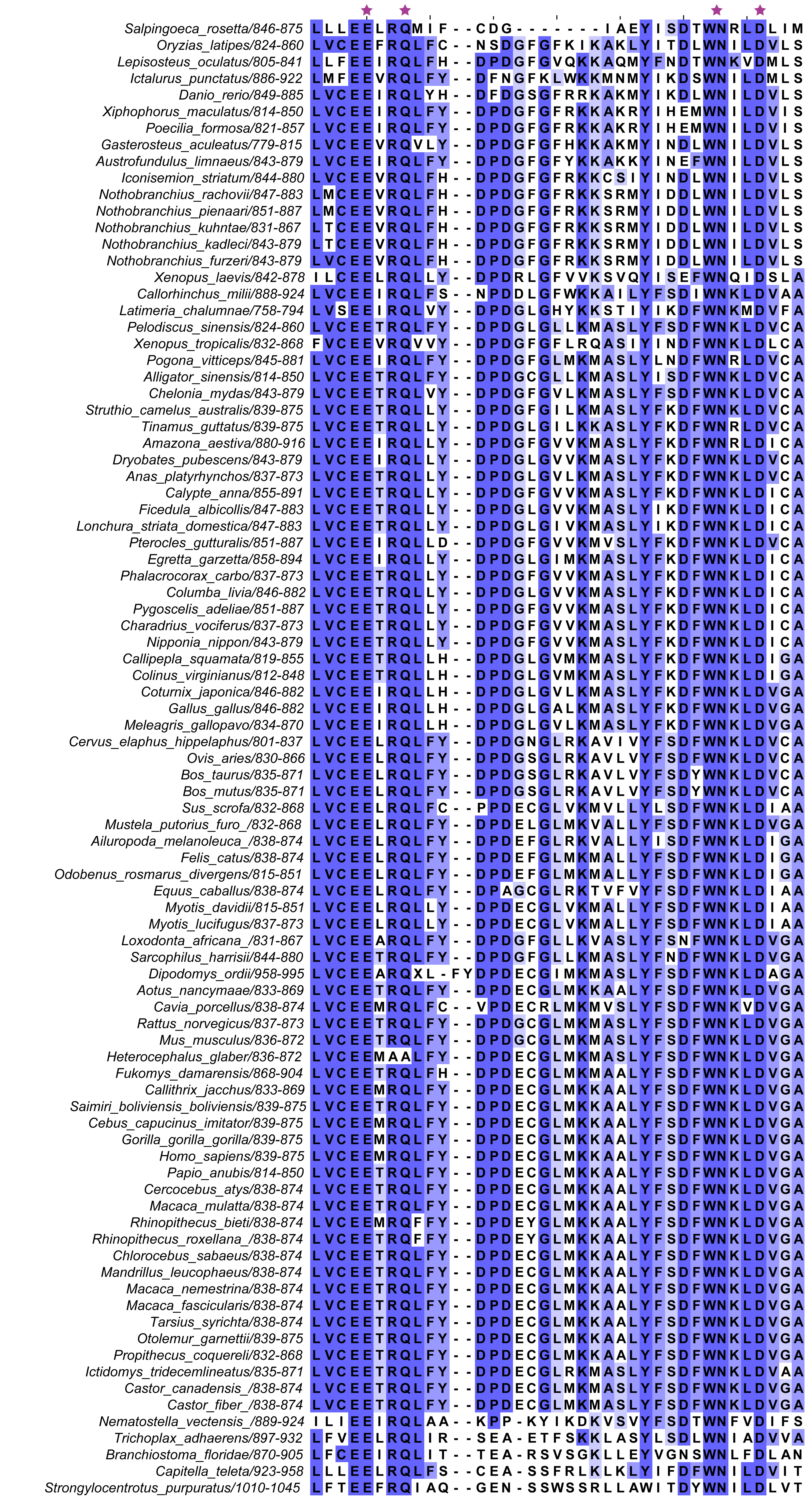

### supplementary Figure S9 89 EF-loop IQ-motif.jpg

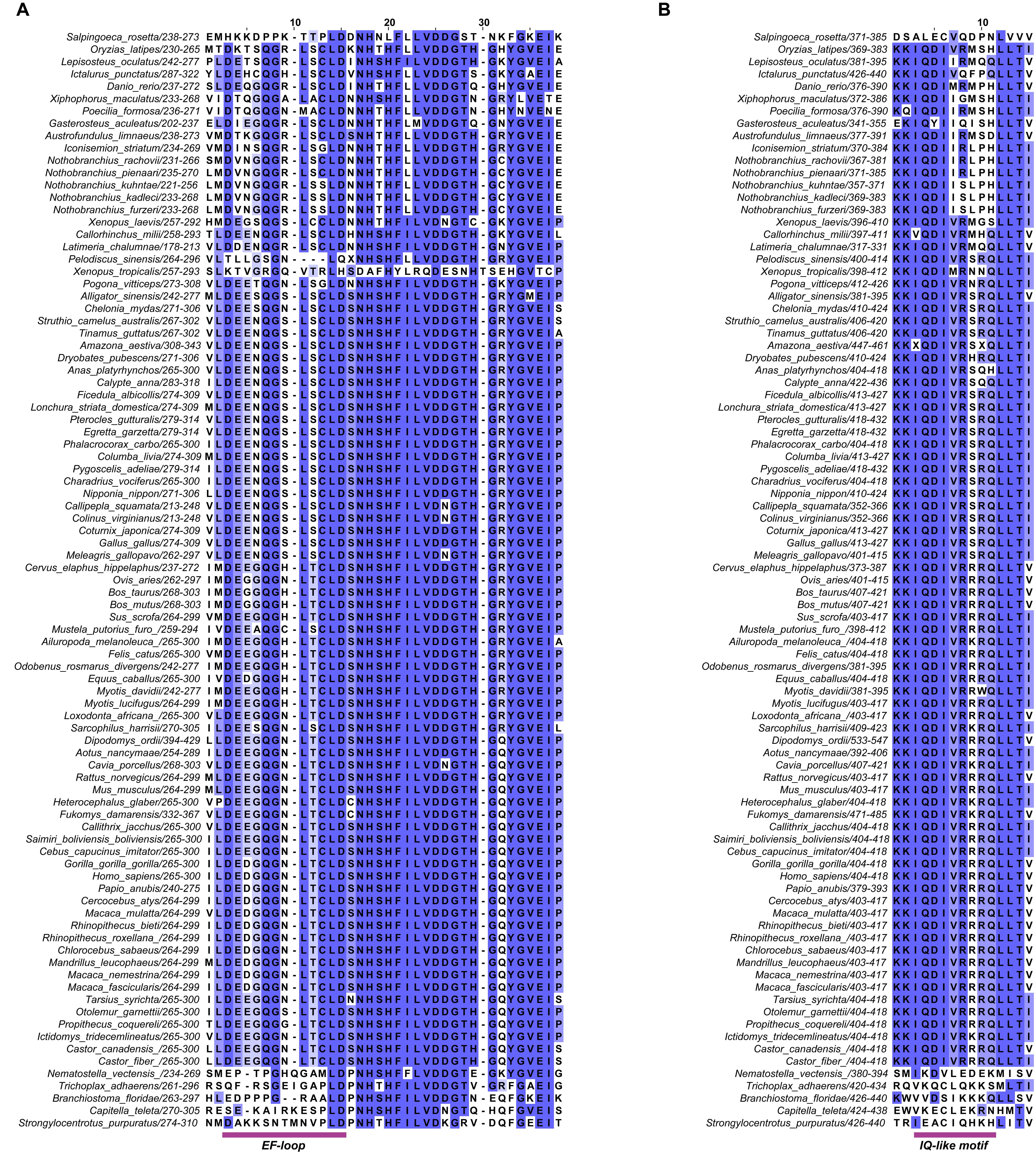
