## Supplementary material for "Evolutionary trajectory of TRPM2 channel activation by adenosine diphosphate ribose and calcium": Table S2 model -2305.docx

| **Model** | **ADPR related channel activation** | | | **Calcium related channel activation** | | | **Requirement of intracellular Ca2+ for its ADPR activation (i.e. Cooperation)** |
| --- | --- | --- | --- | --- | --- | --- | --- |
|  | **N terminal ADPR binding sites in MHR1/2** | **C terminal ADPR binding sites in NUDT9-H domain** | **Chanzyme function** | **Calcium binding sites in S2-S3 helix** | **EF-loop for calcium regulation** | **IQ-like motif for CaM binding** |  |
| **“Primitive” TRPM2** | **Required** | **Not required** | **Yes** | **Required** | **Not required** | **Not required** | **Not required**  **(or only a trace amount)** |
| **“Intermedia” TRPM2** | **Required** | **Not required** | **No** | **Required** | **Not as important** | **Not as important** | **Not required**  **(or only a trace amount)** |
| **“Sophisticated” TRPM2** | **Required** | **Required** | **No** | **Required** | **Required** | **Required** | **Required** |

**Table S2. Summary of activation models for “primitive”, “intermedia” and “sophisticated” TRPM2s.**
